# Sexual dimorphism in histamine regulation of striatal dopamine

**DOI:** 10.1101/2024.05.20.595049

**Authors:** Meghan Van Zandt, Christopher Pittenger

## Abstract

Dopamine modulation of the basal ganglia differs in males and females and is implicated in numerous neuropsychiatric conditions, including some, like Tourette Syndrome (TS) and attention deficit hyperactivity disorder (ADHD), that have marked sex differences in prevalence. Genetic studies in TS and subsequent work in animals suggest that a loss of histamine may contribute to dysregulation of dopamine. Motivated by this, we characterized the modulation of striatal dopamine by histamine, using microdialysis, targeted pharmacology, and shRNA knockdown of histamine receptors. Intracerebroventricular (ICV) histamine reduced striatal dopamine in male mice, replicating previous work. In contrast, and unexpectedly, ICV histamine increased striatal dopamine in females. ICV or targeted infusion of agonists revealed that the effect in males depends on H2R receptors in the substantia nigra pars compacta (SNc). Knockdown of H2R in SNc GABAergic neurons abrogated the effect, identifying these cells as a key locus of histamine’s regulation of dopamine in males. In females, however, H2R had no discernible role; instead, H3R agonists in the striatum increased striatal dopamine. Strikingly, the effect of histamine on dopamine in females was modulated by the estrous cycle, appearing only in estrus/proestrus, when estrogen levels are high. These findings confirm the regulation of striatal dopamine by histamine but identify marked sexual dimorphism in and estrous modulation of this effect. These findings may shed light on the mechanistic underpinnings of sex differences in the striatal circuitry, and in several neuropsychiatric conditions.

**Significance Statement:** Dysregulation of the basal ganglia contributes to the pathophysiology of numerous neuropsychiatric diseases, including several, such as Tourette syndrome (TS), that are characterized by sex differences. Previous genetic studies in humans identified histamine dysregulation as a potential to the development of TS; follow-up work in mice highlighted the role of histamine in regulating striatal dopamine tone. Striatal dopamine dysregulation has been implicated in numerous neuropsychiatric disorders, including TS, schizophrenia, and attention deficit-hyperactivity disorder. Here, we uncover novel and significant sex differences in histamine regulation of dopaminergic modulation of the striatum. A better understanding of these differences may provide insight into the mechanisms underlying sex-dependent outcomes in neuropsychiatric disease.

## Introduction

The basal ganglia are regulated by dopamine (DA) from the midbrain. Striatal DA levels and regulation differ between males and females: Women have been reported to have higher DA synaptic concentration (Laakso et al., 2002), higher DA response to methylphenidate in the ventral striatum (Manza et al., 2022), higher D_2_ receptor levels (Kaasinen et al., 2001; Malén et al., 2022), and higher levels of DA transporter (DAT) (Andersen et al., 2012). The relationship of striatal DA to reward also differs by sex: Men show elevated DA synthesis in response to reward, while women decreased synthesis in response to punishment (Hahn et al., 2021). Many of these effects have been confirmed in animal models, with females showing elevated DA release amplitude (McDermott and Dluzen, 2007; Arvidsson et al., 2014), denser SNc axonal fibers during development (Ovtscharoff et al., 1992), higher DA uptake rates (Walker et al., 2000), and increased accumulation of DA in the absence of DAT clearance (Yu and Liao, 2000; Walker et al., 2006). Sex hormones appear to play a role: estradiol and progesterone potentiate striatal dopaminergic signaling. Ovariectomy significantly decreases DA release and turnover, and both progesterone and estradiol enhance striatal DA responses in females (Becker, 1990; Morissette et al., 1990; Castner et al., 1993; Bazzett and Becker, 1994; McDermott et al., 1994; Becker, 1999; Calipari et al., 2017; Yoest et al., 2018). Furthermore, estrogen receptors are expressed on substantia nigra pars compacta (SNc) dopaminergic neurons that project to the striatum (Creutz and Kritzer, 2004; Munro et al., 2006; Almey et al., 2015).

Many neuropsychiatric disorders differ in epidemiology and presentation between males and females; in some of these, striatal dopamine is implicated in pathophysiology. For example, Tourette syndrome (TS) is diagnosed 3-5 times more often in males than in females (Robertson, 2003; Chou et al., 2022), with sex differences in severity, onset, and comorbidity (Schwabe and Konkol, 1992; Garcia-Delgar et al., 2022; Larsh et al., 2023), and is characterized by increased DA levels in the basal ganglia (Pittenger, 2024). Genetic evidence suggests that some cases of TS are associated with abnormalities in histamine (HA) (Ercan-Sencicek et al., 2010; Pittenger, 2017). Work in animals has focused on the modulation of striatal activity and DA by histaminergic neurons in the tuberomammillary nucleus (TMN) of the hypothalamus. Intracerebroventricular (ICV) infusion of HA reduces striatal DA levels (Castellan Baldan et al., 2014; Pittenger, 2020); notably, these experiments were performed in male mice. Histaminergic neurons project from the TMN throughout the brain, including to the striatum (Takagi et al., 1986; Airaksinen and Panula, 1988) and SNc (Anichtchik et al., 2000). HA binds to four G-protein-coupled receptors (H1R-H4R); the first three of these are expressed in both the striatum and SNc (Honrubia et al., 2000; Haas and Panula, 2003; Zhuang et al., 2018).

Considering the importance of striatal DA to normal basal ganglia function, and its dysregulation in neuropsychiatric disease, the mechanisms by which HA regulates dopamine are of great interest, but they are not well understood. No previous studies have examined the possibility of sex differences in histamine regulation of dopamine. Here we sought to extend our earlier finding that HA reduces striatal DA *in vivo* (Castellan Baldan et al., 2014). We identified an unexpected sexual dimorphism in the effects of HA on DA: it reduces DA levels in males, replicating our previous work, but increases them in females. Mechanistic dissection using specific agonists and cell type-specific receptor knockdown identified H2Rs on GABAergic neurons of the SNc as a key locus of the effect on DA in males, while H3Rs in the striatum are the key regulators in females. Strikingly, the effect in females varies with the estrous cycle. These results identify a new locus of sexually dimorphic effects in striatal DA dynamics. Characterization of sex differences in the neurochemistry and function of the basal ganglia may help elucidate the pathophysiological underpinnings of the marked sex differences seen in many neuropsychiatric conditions.

## Methods

### Animals

All animal use followed protocols approved by Yale’s Institutional Animal Care and Use Committee. Adult male and female wild-type C57BL/6 mice and Gad2-cre mice were group housed in our animal facility in self-ventilating cages, maintained on a 12hr light/dark cycle, and provided with food and water *ad libitum*. Gad2-cre mice were initially obtained from Jackson Laboratories (#010802) then bred in-house. Experiments were performed at 8-10 weeks of age. Separate cohorts of mice were used for each set of experiments due to different cannulation needs. See Figure 1 for a detailed schematic of mice used for each experiment and experimental timelines.

**Figure 1.**
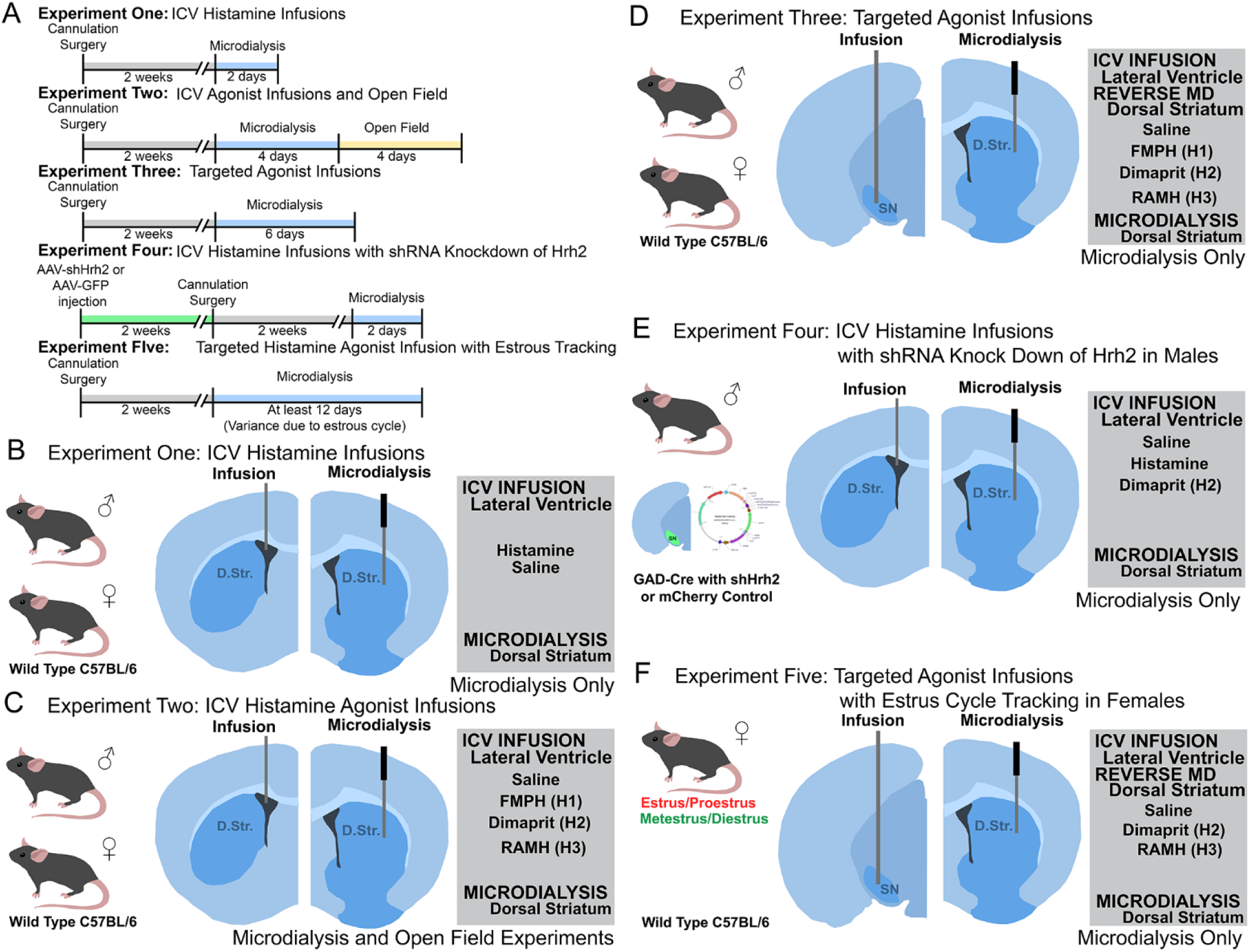
Timeline and methods for experiments. **A.** Timeline of events for each of the five different experimental types described in this manuscript. **B.** Cannula placements and drugs for male and female mice used in ICV Histamine experiments. **C.** Cannula placements and drugs for male and female mice used in ICV Histamine agonist infusion and open field experiments. **D.** Cannula placements and drugs for male and female mice used in Targeted Histamine Agonist experiments. **E.** AAV target, cannula placements, and drugs for male mice used in ICV Histamine Infusion with shRNA knock down of histamine receptor H2 (*Hrh2*) experiments. **F.** Estrus cycle groups, cannula placements, and drugs for female mice used in Targeted Histamine Agonist Infusion with Estrus Cycle Tracking.

### Surgery

For guide cannulation, mice were anaesthetized using a combination of ketamine/xylazine (10mg/kg / 1mg/kg) until completely sedated as measured by lack of response to noxious stimuli. The skull was exposed, and an anchor screw was inserted between the two sites of cannula implantation for implant stability. Two holes were drilled, one for each cannula. Microdialysis guide canulae (CMA7) were implanted at AP+0.9 and ML+/-1.5 at a depth of 1.0mm to allow the 2mm length microdialysis membrane when inserted to enter directly into the dorsal striatum (Figure 1, 2A). Infusion guides were implanted at one of two locations, depending on the experiment. ICV guide cannulae were implanted at AP+0.4, ML+/-0.7, and DV+2.3, contralateral to the microdialysis cannula, to target the lateral ventricle (Figure 1B, 2B). Guide cannulae for SN-targeted infusion were implanted at AP-3.0, ML+/-1.5, and DV+4.5, ipsilateral to the microdialysis guide (Figure 1D, 3D). Guide cannulae were cemented in place using Metabond Dental cement (Parkell). Following surgery, mice recovered for at least one week prior to microdialysis experiments. See Figure 1B-F for cannula placements.

For short hairpin RNA (shRNA) experiments (Figure 1E), we used an adeno associated virus (AAV) generated by VectorBuilder expressing a shRNA against the murine *Hrh2* gene and an eGFP reporter within a cre-activated DIO cassette (Vector ID: VB200728-1049hdn) for controlled expression in cells expressing Cre. GAD-cre mice were anesthetized as described above, the skull exposed, and holes drilled bilaterally at AP-3.0, ML+/-1.5. A Hamilton syringe (1.0µl, 32ga, blunt) containing either the shRNA virus or a scramble GFP control was lowered to DV+4.5 and 0.5µl of virus was injected slowly over three minutes. The needle remained in place for two minutes following injection to prevent aspiration of the virus as the needle is removed. Following AAV injection, the scalp was sealed by VetBond and mice recovered for two weeks prior to proceeding to canulation surgery, as detailed above. Viral targeting and reduction of H2 were confirmed following experiments using immunohistochemistry (see Figure 4C).

### Microdialysis

Mice were acclimated to the testing facility for one hour on the day prior to microdialysis. On test days, mice were habituated to the microdialysis testing facility for 30 minutes prior to any handling. For estrous cycle testing, female mice were restrained prior to the acclimation period and 10µl of sterile saline was injected into the vaginal cavity using a blunt pipette tip, then quickly aspirated. Vaginal samples were applied to a glass slide and visualized on a light microscope (AmScope). Estrous cycle stage was identified by cell density and composition, as previously described (Byers et al., 2012). All mice were handled before experimentation to acclimate them. Restraint was performed on males in a similar manner prior to experimentation to reduce variability in handling.

All microdialysis experiments were performed in the CMA120 System for freely moving animals (Harvard Apparatus; protocol modified from Castellan Baldan et al, 2014). Infusion of ACSF (Harvard Apparatus, #597316) was performed using a CMA402 Microdialysis Syringe pump set at a rate of 2µl/min. At the start of the experiment, mice were briefly restrained, the dummy protective cannula was removed from the guide, and a CMA7 microdialysis probe (6 kDa, 2mm membrane length, Harvard Apparatus) was gently inserted. Mice were then placed into the apparatus and flow was turned on. ACSF was allowed to flow through the tubing for 30 minutes to fully account for any delay in sample travel to collection, followed by another 30 minutes of acclimation. Following acclimation, samples were collected every ten minutes and put immediately on dry ice. After one hour of baseline sample collection, mice were given drug or saline, and samples were collected for another 90 minutes. Mice received one treatment per day on consecutive days; drug treatment order was randomized for each mouse, and randomization order was included as a covariate in all analyses. Some mice were unable to complete all drug treatments due to cannula malfunction; for this reason the N differs in some treatment groups.

For infusion of drug (ICV and SN), mice were briefly restrained and an infusion cannula matched to the length of the guide (2.3mm for ICV, 4.5mm for SN infusion) was inserted into the guide. Drugs were infused at a rate of 1µl/min for 2 minutes total. The infusion cannula was left in place for one minute following the end of drug infusion, to prevent backflow. For drug infusion into the striatum, reverse microdialysis was used; drug was infused through the microdialysis cannula at double the concentration used for direct infusions. Mice were briefly restrained prior to the initiation of reverse microdialysis infusion, such that the effects of restraint stress prior to drug were matched between groups.

### Drugs

Drug doses were: HA 20µg/mouse (Castellan Baldan et al., 2014), 2-(3-trifluoromethylphenyl)histamine (FMPH) 6µg/mouse (Malmberg-Aiello et al., 1998), Dimaprit 20µg/mouse (Michinaga et al., 2022), and RAMH 20µg/mouse (Vohora et al., 2001; Rapanelli et al., 2017b), prepared in sterile saline. All agonists were chosen for specificity for their receptor (Durant et al., 1977; Arrang et al., 1987; Krause et al., 1995; Malmberg-Aiello et al., 1998) and all doses were chosen based on literature review based on previous ICV or brain infusion without toxicity in mice (Oishi et al., 1989; Vohora et al., 2001; Purón-Sierra and Miranda, 2014; Rapanelli et al., 2017a; Patel et al., 2021; Michinaga et al., 2022). Drug treatments were given in a randomized order per animal.

### HPLC

Mouse striatal microdialysis samples were snap-frozen on dry ice and stored at −80° prior to quantification in batches using HPLC. DA concentration was measured on a DIONEX UltiMate3000 RS Electrochemical Detector (Thermo-Fisher) using a computer running Chromeleon 7 software (Thermo-Fisher). Samples were run through a Hypersil ODS C18 column with a 3µm particle size, 3mm diameter and 150mm length (Thermo-Fisher) and analyzed using a 6011RS ultra-analytical cell (Thermo-Fisher). Samples were run using MDTM mobile phase (Thermo-Fisher) at a rate of 0.5mL/min through the column and analyzed using electrode potentials at 500nm/-500nm with a 0.2s filter over a run time of 10 minutes. Using these parameters, DA standards reliably produced a peak even at low concentrations between 3:45 and 4:15 after injection; as such, peaks in our samples were analyzed within this time window. Once samples were run, we checked for abnormal background or false peaks (which can be caused by air bubbles in the sample); samples were rerun if anomalies were apparent. HPLC data was analyzed using Cobra Wizard (Thermo-Fisher) to measure peak heights against baseline. Dopamine is represented as % change from baseline over 20-60 min post-infusion, to control for between-animal and between-session variance. Any sample sets in which a stable baseline was not able to be established were not included in data analysis.

### Behavior

Open field (OF) recordings were performed using the Fusion Sensor System and Experimentation Software (Omnitech Electronics, Inc.). Following 30 minutes acclimation to the behavioral room, mice were placed in the open field apparatus (16×16in) and baseline data collected for one hour. After one hour of baseline, mice were briefly returned to their home cage and administered drug as described above. Mice received one treatment per day, in a counterbalanced order. Following drug administration, mice were returned to the OF and their total activity was measured for another hour. All OF data was normalized to the same day’s baseline to control for individual animal and day effects.

### Immunohistochemistry

For histology, mice were anesthetized (ketamine 100mg/kg and xylazine 10mg/kg, followed by isoflurane inhalation until no response to noxious stimulus) followed by transcardial perfusion with 4% paraformaldehyde in phosphate buffered saline (PBS) (Thermo-Fisher) and brain extraction. Cryostat sections were cut at 30µm and mounted on Diamond White glass slides (Globe). For immunostaining, slides were incubated overnight at room temperature with primary antibodies diluted in 0.1 M PBS containing 10% normal donkey serum (Abcam, ab7475) and 1% Triton-X 100 (AmericanBio). Primary antibodies were Rabbit anti-GFP (1:1000, Thermo-Fisher, A-11122) and goat anti-H2R (1:250, Everest Biotech, EB06905). Following incubation with primary antibody, slides were washed three times for ten minutes each in PBS with 0.1% Triton at room temperature. Detection was performed with appropriate secondary antibodies, again in PBS + 10% normal donkey serum and 1% Triton: polyclonal donkey anti-rabbit coupled to Alexa Fluor 488 (1:1000, Thermo-Fisher, A-21206) and polyclonal donkey anti-goat coupled to Alexa Fluor 568 (1:1000, Thermo-Fisher, A-11057). Following a two-hour incubation in secondary antibody, slides were rinsed three times at twenty minutes each in PBS and then mounted and coverslipped in ProLong Gold with DAPI (Thermo-Fisher, P36935). Immunostained coronal SNc sections (30 µm), identified following landmarks in the atlas of Paxinos (Franklin and Paxinos, 1997), were visualized on a Zeiss Scope.A1 using a Plan-APOCHROMAT 10x objective and an Axiocam 503 Mono at 100x magnification.

### Quantification of H2R knockdown

Images were imported into FIJI ImageJ (Schindelin et al., 2012) for analysis using the plugin Puncta Analyzer (Savage et al., 2024) for unbiased quantification. Using this plugin, we quantified the number of red fluorescent H2R puncta colocalized with GFP-expressing GAD-cre GABAergic neurons. During quantification, thresholds were set at the base of the histogram, typically at the value recommended by the program, so that background fluorescence would be excluded during analysis. A total of 119 neurons from 5 GAD-Cre GFP controls and 245 neurons from 5 GAD2-shH2 knock down animals were analyzed to confirm efficacy of virus.

### Statistical analysis

All data were imported into Graph Pad Prism 10 (Graph Pad Software, LLC) or SPSS (IBM) for statistical analysis. The following analyses were used for each experiment:

**Table One.** Statistical Analyses.

| Experiment | Groups | Treatments | Statistical Test |
| --- | --- | --- | --- |
| ICV Histamine (Over Time) | 2 (males vs females) | 2 (saline vs histamine) | 2-Way Repeated Measures ANOVA (RM-ANOVA) (sex x treatment) |
| ICV Histamine (Collapsed) | 2 (males vs females) | 2 (saline vs histamine) | 2-Way ANOVA (sex x treatment)<br>t-test (within sex comparisons) |
| ICV Infusions (Over Time)<br>Males | 1 (males) | 4 (saline, H1, H2, H3 Agonists) | Repeated Measures ANOVA (RM-ANOVA) |
| ICV Infusions (Over Time)<br>Females | 1 (females) | 4 (saline, H1, H2, H3 Agonists) | Repeated Measures ANOVA (RM-ANOVA) |
| ICV Infusions (Collapsed) | 2 (males vs females) | 4 (saline, H1, H2, H3 Agonists) | Linear Mixed Model (Accounting for Order Effects) |
| TG Infusions (Over Time)<br>Males | 1 (males) | 6 (Striatum – Saline, H2, H3, SN – Saline, H2, H3) | Repeated Measures ANOVA (RM-ANOVA) (brain region x treatment) |
| TG Infusions (Over Time)<br>Females | 1 (females) | 6 (Striatum – Saline, H2, H3, SN – Saline, H2, H3) | Repeated Measures ANOVA (RM-ANOVA) (brain region x treatment) |
| TG Infusions (Collapsed)<br>Males | 1 (males) | 6 (Striatum – Saline, H2, H3, SN – Saline, H2, H3) | Linear Mixed Model with 2-Way Comparison (Accounting for order effects)(brain region x treatment) |
| TG Infusions (Collapsed)<br>Females | 1 (females) | 6 (Striatum – Saline, H2, H3, SN – Saline, H2, H3) | Linear Mixed Model with 2-Way Comparison (Accounting for order effects)(brain region x treatment) |
| shRNA H2 with ICV Infusion (Over Time) | 2 (GAD-cre Control Males, GAD-cre shH2 males) | 2 (saline vs histamine) | Repeated Measures ANOVA (RM-ANOVA) |
| shRNA H2 with ICV Infusion (Collapsed) | 2 (GAD-cre Control Males, GAD-cre shH2 males) | 2 (saline vs histamine) | 2-Way ANOVA (virus x treatment) |
| Estrous TG Infusions (Over Time) | 2 (Met/Di Females, Pro/Est Females) | 4 (Striatum Saline, Striatum H2, SN Saline, SN H2) | 3-Way RM-ANOVA |
| Estrous TG Infusions (Collapsed) | 2 (Met/Di Females, Pro/Est Females) | 4 (Striatum Saline, Striatum H2, SN Saline, SN H2) | Linear Mixed Model with 3-Way Comparison (Accounting for order effects)(estrous phase x brain region x treatment) |
| Open Field with ICV Infusion | 2 (males and females) | 4 (Saline, H1, H2, H3) | 2-Way ANOVA (sex x treatment)<br>ANOVA (within sex comparisons) |

## Results

### Experiment One: ICV Histamine Infusions. Histamine regulates striatal DA levels differently in male and female mice

We previously showed that HA reduces DA levels in the striatum (Castellan Baldan et al., 2014; Rapanelli et al., 2014), but those experiments were limited to males. We cannulated adult male and female mice and infused HA or saline ICV while measuring striatal DA using microdialysis (Figure 1B, 2A,B). Direct comparison of HA’s effects on DA between the sexes revealed significant differences (Figure 2C. two-way RM-ANOVA: sex x treatment F[1,40]=6.799, p=0.013; sex F[1,40]=4.429, p=0.042). As histamine and histamine agonists are quick acting when infused directly into the brain, we did not expect to see an effect of time in our data post infusion; this is consistent throughout all experiments (Table 2). To simplify analysis and visualization, we collapsed data over 20-60 minutes; this yielded similar effects (Figure 2D. two-way ANOVA: sex x treatment, F[1,49]=11.61, p=0.001, main effect of sex, F[1,49]=6.281, p=0.016, trend effect of treatment, F[1, 49]=3.317, p=0.074). Having seen significant sex differences, we therefore analyzed the effect of HA on striatal DA separately in male and female mice. Infusion of ICV HA into male mice replicated previous findings – striatal DA levels 20-60 minutes post-infusion were significantly decreased compared to saline controls both over time (Figure 2C; RM-ANOVA: main effect of treatment, F[1,13]=5.668, p=0.033) and when collapsed across all time points (Figure 2D; t-test: t[29]=2.961, p=0.006). Males showed a slight elevation of DA following saline infusion, likely as a result of the stress of handling and infusion.

**Figure 2.**
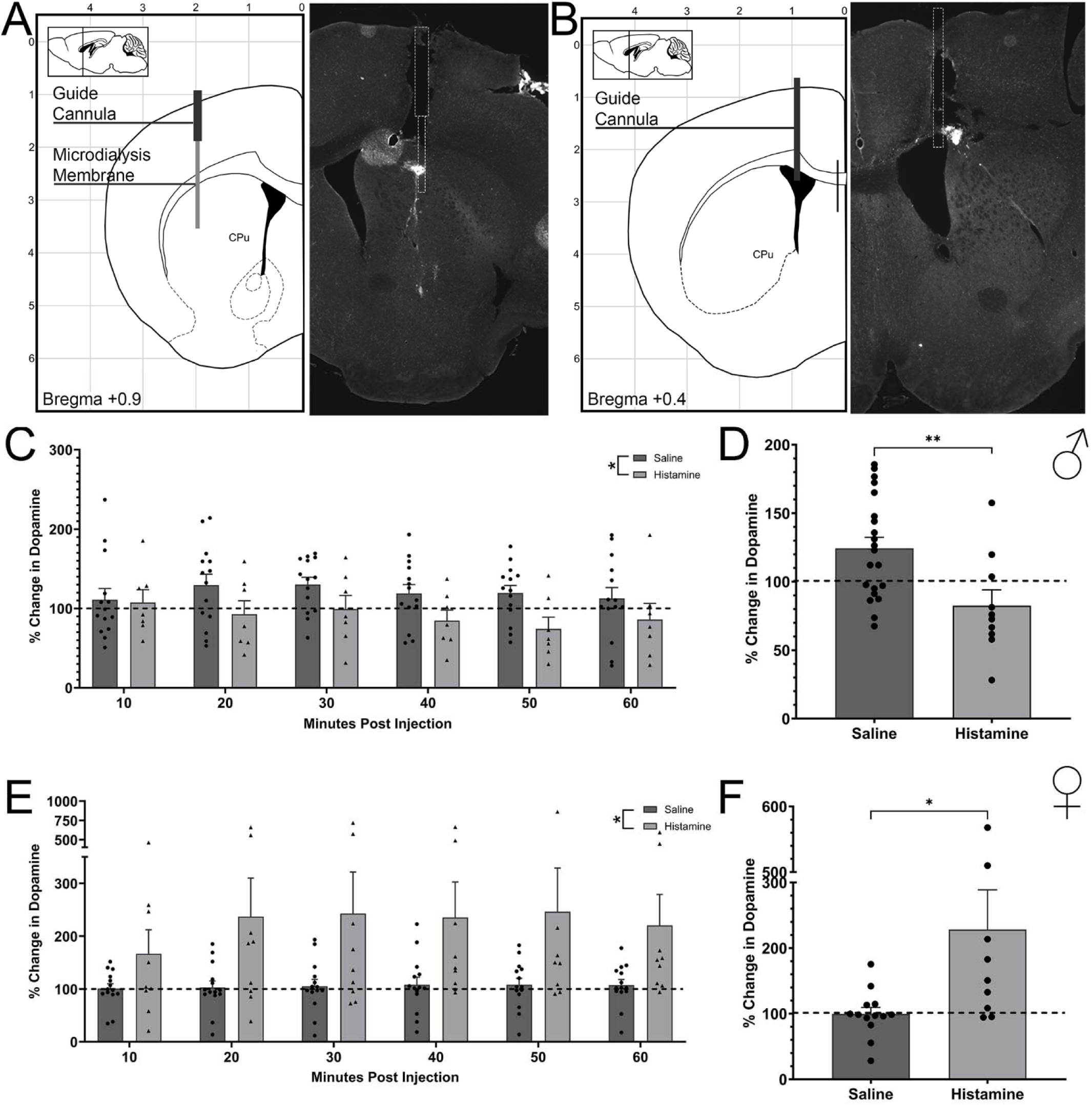
Histamine regulates striatal dopamine differently in males and females. **A.** Schematic and representative image for placement of the microdialysis guide cannula in the striatum. Guide cannula placement was counterbalanced between left and right striatum. **B.** Schematic and representative image for guide cannula placement for ICV infusion. ICV guide cannulae were placed contralaterally to microdialysis guide cannulae. **C.** ICV infusion of HA significantly decreases striatal DA, compared to saline control (sal), in male mice (RM-ANOVA: main effect of treatment, F[1,13]=5.668, *p=0.033). **D.** Combining data over the 20-60 minute interval post infusion confirmed a significant decrease in striatal DA after HA relative to saline (Student’s t-test (two-tailed): T[29]=2.961, **p=0.006). **E.** ICV infusion of HA significantly increased striatal DA, relative to saline-infused controls, in female mice (RM-ANOVA: main effect of drug, F [1,13]=5.633, *p=0.034). **F.** This was confirmed when data were collapsed over 20-60 min post -infusion (Shapiro-Wilk test for Normality, Histamine W=0.734, p=0.004 [no pass], Mann-Whitney: *p=0.024). Dashed lines indicate baseline dopamine levels, determined for each animal during the 50 min before HA or saline infusion.

**Table Two.**
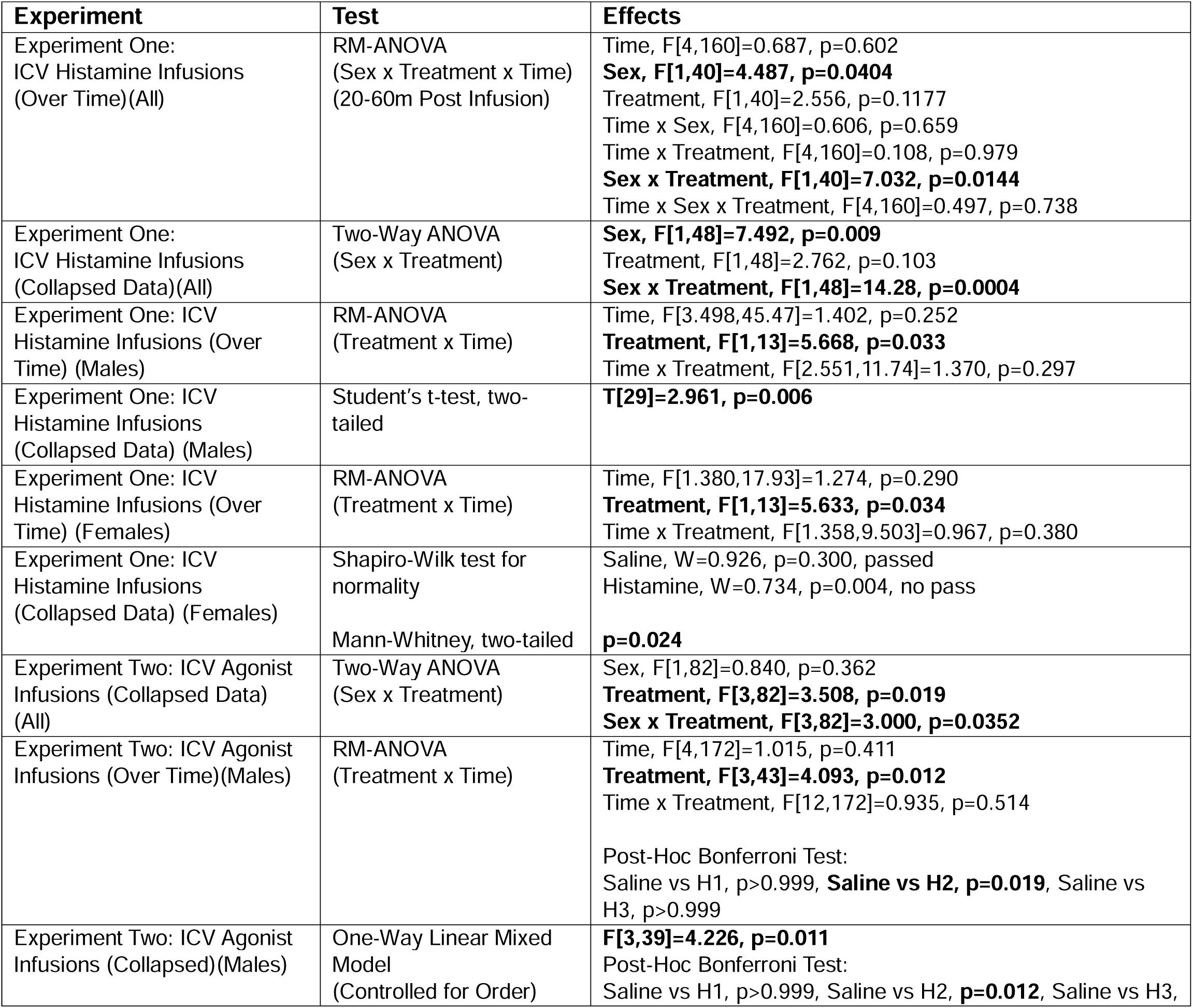
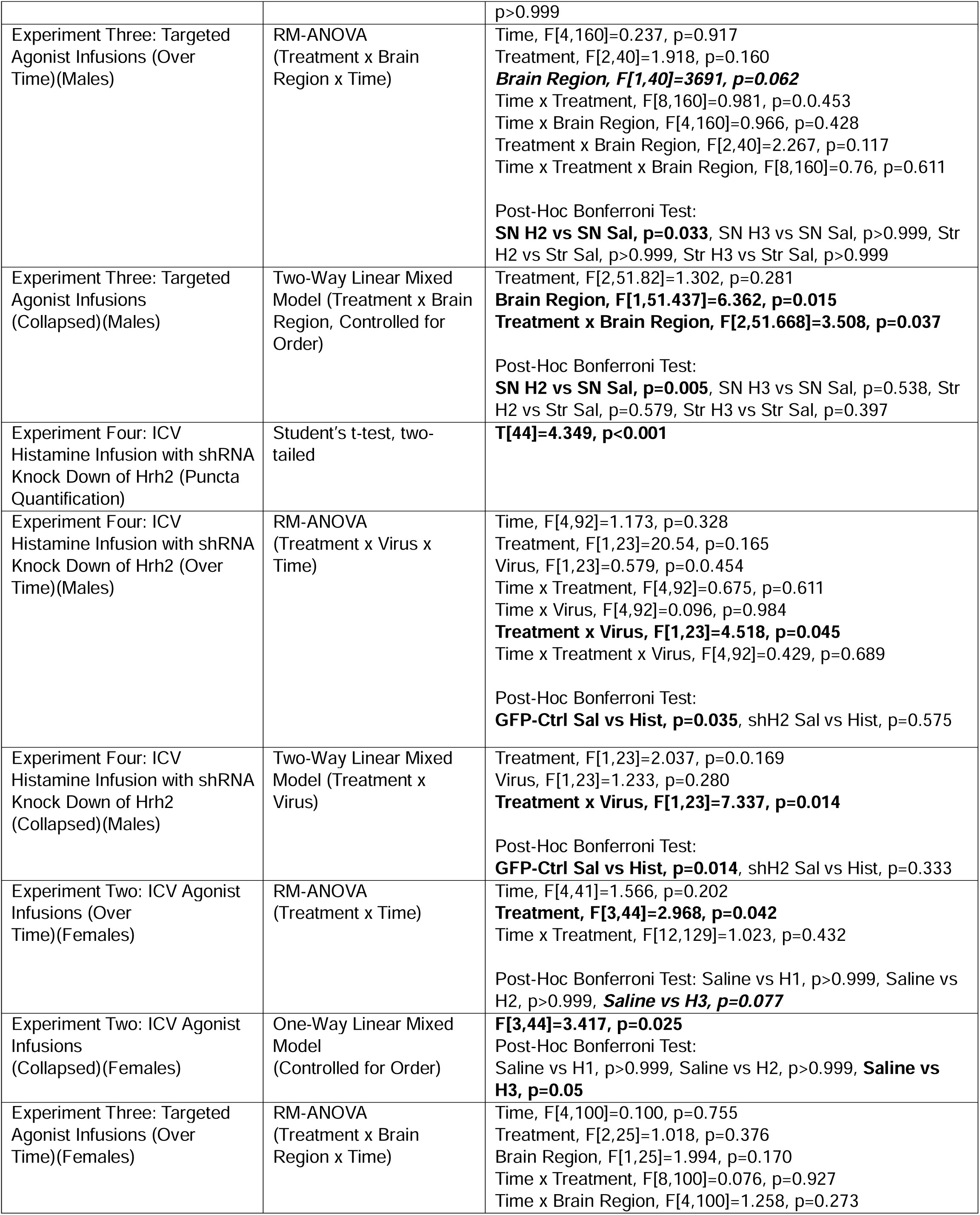
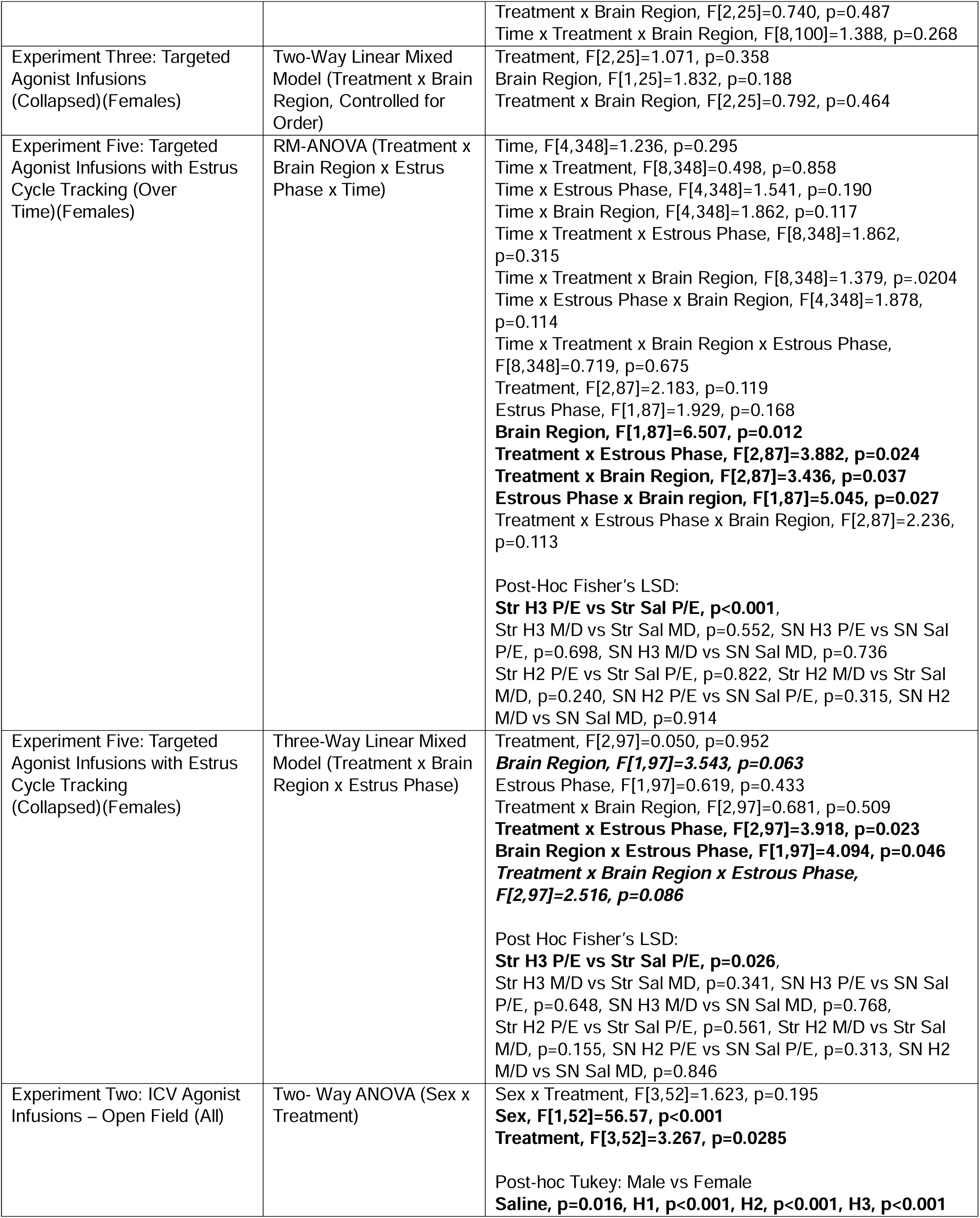
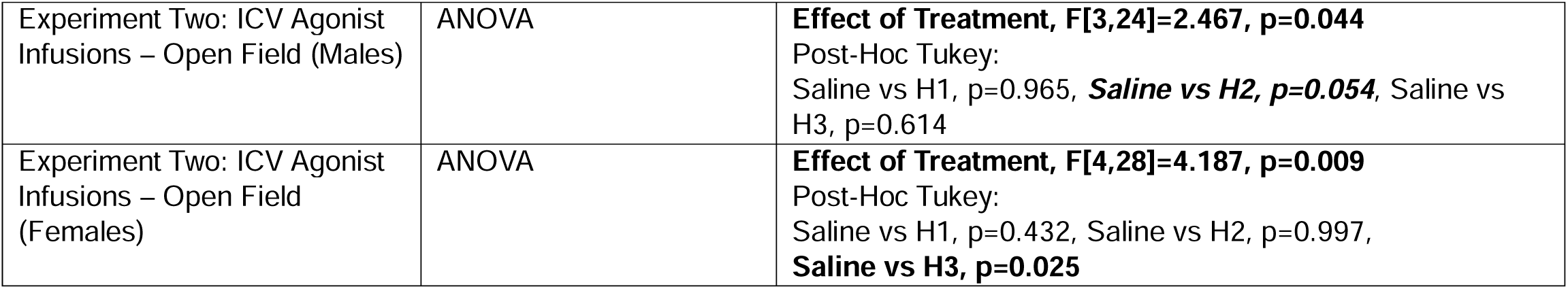
Full Statistical Analyses from All Experiments.

Surprisingly, females showed a very different pattern: DA was significantly increased after ICV infusion of HA, both over time (Figure 2E; RM-ANOVA: main effect of treatment, F[1.000, 13.00]=5.633, p=0.034) and collapsed over the full interval (Figure 2F; Mann-Whitney: p=0.024). Notably, the female data showed a high variance, with a small number of mice with very high DA levels following HA infusion appearing to drive the effect; we return to this point below. (This deviation from normality, which was supported by a significant Shapiro-Wilk test, motivates the use of a non-parametric test in Figure 2F). Saline-treated females showed no elevation in DA. In sum, these results show significantly different effects of histamine on striatal dopamine in males and females – histamine infusion decreases striatal dopamine in males but increases it in females.

### Experiment Two: ICV and Targeted Histamine Agonist Infusion. H2Rs suppress striatal DA in males and decrease locomotion

To determine which HA receptor regulates the effects of HA on striatal DA, we infused specific HA receptor agonists intracerebroventricularly (ICV)(Figure 1C). Once again, direct comparison of agonist effects on striatal DA indicates significant differences between sexes (two-way ANOVA: sex x drug, F[3,82]=3.000, p=0.035, main effect of drug, F[3,82]=3.508, p=0.019); we therefore analyzed effects in males and females separately. Analysis of change in DA over the 20-60 minutes following infusion in males indicates a significant effect of treatment (Figure3A; RM-ANOVA, effect of treatment, F[3,43]=4.093, p=0.012), and post-hoc analysis showed that infusion of the H2 agonist dimaprit was the only treatment that significantly decreased DA levels from baseline, relative to saline (Post-hoc Bonferroni: Saline vs H2 Agonist p=0.019, Saline vs H1 p>0.999, Saline vs H3 p>0.999). We again collapsed this data over 20-60 minutes; since animals received repeated infusions in counterbalanced order in this experiment, we used LMEM controlling for order for this analysis. This showed the same H2 effect (Figure 3B; LMEM: effect of treatment, F[3,39]=4.226, p=0.011, Bonferroni Post-hoc Saline vs H2 p=0.012, other pairwise comparisons with saline N.S.). These data indicate that the H2 receptor in males mediates the effects of HA on DA, while H1 and H3 have no effect.

**Figure 3.**
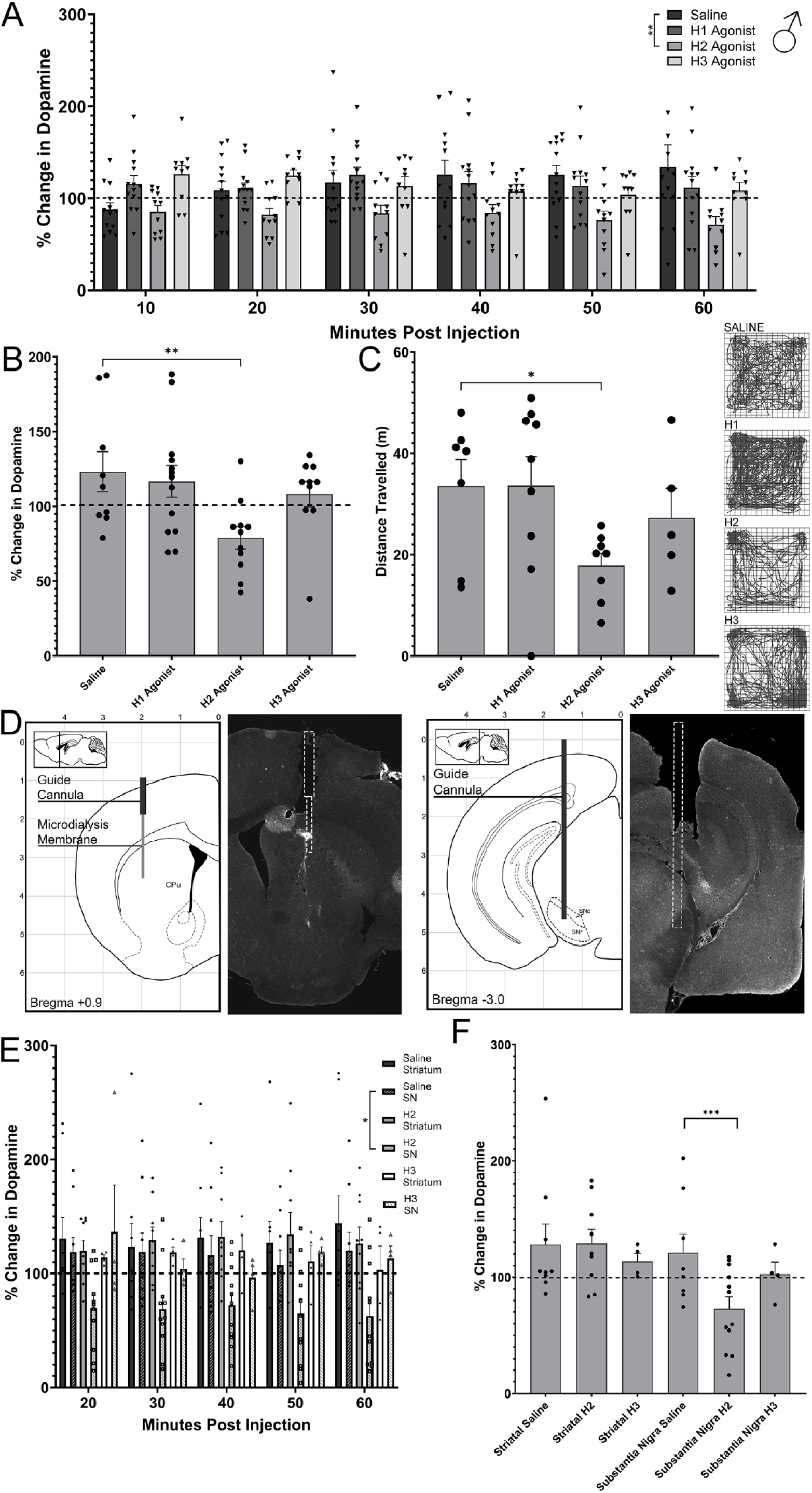
Striatal dopamine is regulated by activation of histamine H2 receptors in the SNc of male mice. **A.** ICV infusion of HA H2R agonist dimaprit is the only treatment to reduce striatal dopamine over the 20-60 minute post drug infusion interval (RM-ANOVA: Effect of Treatment, F[3,43]=4.093, p=0.012, Post-Hoc Bonferroni **Saline vs Dimparit (H2) p=0.019, Saline vs FMPH (H1), p>0.999, Saline vs RAMH (H3) p>0.999). **B.** When collapsed over the 20-60 minute time period, ICV infusion of the HA H2R agonist dimaprit significantly reduces striatal DA in male mice, compared to saline controls and all other agonist treatments. (LMEM: Effect of Treatment, F[3, 36]=3.625, p=0.022; Post-Hoc Bonferroni test: Sal vs Dimaprit **p=0.032). No effect of the H3R agonist RAMH or H1R agonist FMPH was seen (Bonferroni’s’s (sal vs RAMH): p>0.99, (sal vs FMPH): p>0.999) **C.** Locomotion in male mice following treatment with HA agonists was reduced by dimaprit relative to saline control (ANOVA: Effect of Treatment, F[3, 24]=2.467, p=0.044; Tukey’s (sal vs dimaprit): p=0.054). Representative traces for the open field are shown to the right. **D.** Schematic and representative image for guide cannula placement of microdialysis guide cannula in the striatum and infusion guide cannula above the substantia nigra; drugs targeting the striatum in this experiment were delivered through the same cannula using reverse microdialysis. Guide placement was counterbalanced between left and right striatum. SN infusion guides were placed ipsilateral to microdialysis cannula. **E.** Infusion of histamine agonists into either the striatum or SNc shows a strong trend towards effect of brain region over time, with post-hoc testing indicating that infusion of dimaprit H2 agonist into the SNc is the only treatment to significantly reduce dopamine (RM-ANOVA: Brain Region, F[2,51]=3.691 p=0.062, Post-Hoc Bonferroni: *SN Sal vs SN H2 p=0.033, SN H2 vs Str H2 p=0.002, all other treatments/regions N.S.) **F.** When collapsed over the 20-60 minute post infusion interval, infusion of dimaprit into the SN significantly reduces DA levels in the striatum, compared to saline controls and the H3R agonist RAMH (LMEM: Interaction Treatment x Brain Region F[6, 51]=3.508, p=0.037, effect of Brain Region F[2,51]=6.361, p=0.015, Bonferroni (SN Sal vs SN H2 p=0.005, Str Sal vs Str H2 p=0.579, SN H2 vs Str H2 p<0.001, SN Sal vs SN H3 p=0.538, Str Sal vs Str H3 p=0.397).

As ICV HA infusion has previously been shown to cause hypolocomotion (Castellan Baldan et al., 2014), which may be associated with its effects on DA, we tested male mice in the open field after HA receptor agonist infusion. In males, ICV infusion of the H2R agonist dimaprit produced a trend towards decreased locomotion (Figure 3C; 1-way ANOVA: F[3, 24]=3.132, p=0.044; post-hoc Tukey’s saline vs H2, p=0.054; other pairwise comparisons with saline N.S.).

### Experiment Three: Targeted Histamine Agonist Infusion. H2R activation suppresses striatal DA in males

To determine the locus of the H2R effect in males, we infused the H2 agonist dimaprit into both the striatum and the SN (Figure 1D, 3D). Based on ICV results (Figure 3A,B) we did not expect an effect of H3 agonist activation; this was confirmed in a few mice infused with the H3 agonist RAMH. The over-time data reveals a trend-level effect of brain region, with significant differences when H2 is activated in the SNc (Figure 3E; RM-ANOVA: Effect of Brain Region, F[1,40]=7.131, p=0.062, Post-hoc Bonferroni: SN Saline vs SN H2, p=0.033; other pairwise comparisons for region or H3 N.S.). Collapsed over time, there was a significant effect of brain region (Figure 3F; LMEM: F[1,51]=6.362, p=0.015) and a significant interaction of brain region and treatment (F[2,52]=3.508, p=0.037), with decreased striatal DA following infusion of dimaprit into the SNc, relative to saline (Post-hoc Bonferroni: p=0.005). Targeted infusion of dimaprit into the striatum or of RAMH into either the SNc or the striatum had no effect (Striatum H2 vs Saline, p=0.579, Striatum H3 vs Saline, p=0.379, SN H3 vs Saline, p=0.178). Therefore, we conclude that activation of H2 receptors in the SN in males reduces striatal dopamine levels.

### Experiment Four: Short Hairpin RNA Knock Down of H2Rs in Males with ICV Histamine Infusion. H2Rs on GABAergic interneurons in the SNc mediate the H2-induced decrease of striatal DA in males

As the Gαs-coupled H2R typically has a net excitatory effect on neurons, we hypothesized that it reduces striatal DA indirectly by activating SNc GABAergic interneurons. To test this, we specifically downregulated H2R in SNc GABAergic interneurons by infusing recombinant AAV expressing an anti-H2R shRNA in a cre-dependent fashion into the SNc of male GAD2-cre transgenic mice (Figure 1E, 4A-C). Receptor knockdown was confirmed using immunohistochemistry with cell-specific quantification (Figure 4C, Student’s t-test, t[44]=4.349 p<0.001). ICV HA infusion into these mice once again reduced striatal DA in GAD2-cre mice receiving a negative control GFP-expressing virus, but this effect was absent in mice in which H2R was knocked down. This effect was consistent when examined over time (Figure 4D; 3-way RM-ANOVA: virus x treatment, F[1,23]=4.518, p=0.044, post-hoc Bonferroni GFP-Ctrl Sal vs Hist, p=0.035, shH2 Sal vs Hist, p=0.575) and when collapsed over the 20-60 minute post injection interval (Figure 4E; two-way ANOVA: virus x treatment, F[1,23]=7.337, p=0.014, effect of virus, F[1,23]=1.233, p=0.280, effect of treatment, F[1,23]=2.037, p=0.169). ICV HA infusion significantly decreased DA, relative to saline infusion, in GAD2-Cre-GFP controls (post-hoc Bonferroni: GAD2-Cre-GFP p=0.014) but not in GAD-cre shH2 mice (p=0.333). Similar effects were seen after ICV infusion of the H2 agonist dimaprit (Figure 4F; 2-way ANOVA: interaction virus x treatment, F[1,11]=7.042, p=0.022; Tukey’s: GFP, p=0.044; shH2, p=0.189). GAD2-Cre mice receiving GFP control virus did not differ significantly from wild type controls in any experiment (p>0.05). As such, we conclude that the H2 receptors on SN GABAergic inhibitory interneurons are mediating the decrease in striatal dopamine in males.

**Figure 4.**
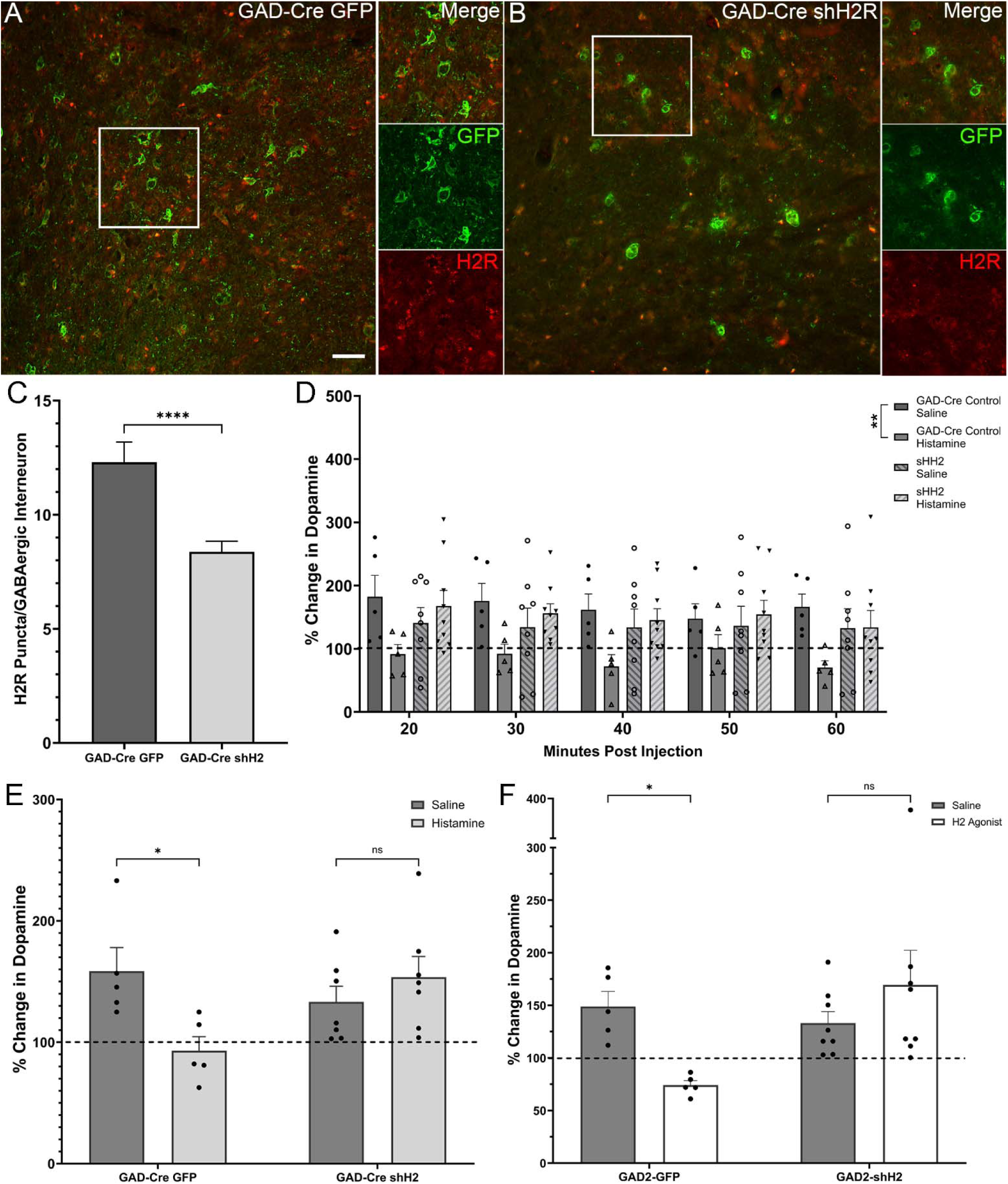
GABAergic interneurons in the substantia nigra mediate the effects of H2 activation on striatal DA in males. **A, B.** Representative image of GFP expressing GABAergic interneurons (green) and H2R (red) in the SN of a mouse that received control scramble shRNA AAV (A) or AAV-shH2 (B). Higher resolution images of selected regions demonstrate reduced expression of H2R on GFP-expressing GABAergic interneurons. Scalebars: 20µm. **C.** Quantification of colocalization of red fluorescent H2R puncta with GFP-expressing GABAergic interneurons shows a significant decrease in the quantity of colocalized H2R in mice that received the shH2 AAV compared to controls (Student’s t-test (two-tailed), T[44]=4.349, ****p<0.001). **D.** H2R knockdown in GABAergic interneurons in the SN blocks the decrease of striatal dopamine by histamine ICV infusion over the 20-60 minute time interval post injection (RM-ANOVA: Interaction Virus x Treatment, F[4,23]=4.518, p=0.045, Post-Hoc Bonferroni: Gad-Cre GFP CTRL (sal vs HA) p=0.035, Gad-Cre shH2 (sal vs HA) p=0.575). **E.** H2R knockdown in GABAergic interneurons in the SN blocks the reduction of striatal DA after ICV HA infusion [2-Way ANOVA: interaction Virus x Treatment: F[1, 23]=7.337 p=0.014; Post-Hoc Bonferroni: Gad-Cre GFP CTRL (sal vs HA), p=0.014, Gad-Cre shH2 (saline vs HA), p=0. 333, HA (GFP vs shH2) p=0.014]. **F.** H2R knockdown in GABAergic interneurons in the SN blocks the effects of ICV dimaprit infusion on striatal DA levels (2 Way ANOVA: interaction Virus x Treatment: F[1, 11] = 4.068, p=0.022; Tukey’s: Gad-Cre GFP CTRL (sal vs dimaprit): *p=0.044, shH2 (sal vs dimaprit): p=0.189). Dashed lines indicate baseline striatal DA levels, prior to drug vs saline infusion.

### Experiment Two: ICV Histamine Agonist Infusion in Females. H3R activation increases striatal DA in females

We repeated this analysis in female mice, with strikingly different results. ICV infusion of HA receptor agonists in females (Figure 1C) again showed significant effects 20-60 minute post-treatment, both over time (Figure 5A; RM-ANOVA: effect of treatment, F[3,44]=2.968, p=0.042) and when collapsed over the interval (Figure 5B; LMEM: F[3,44]=3.417, p=0.025). Here, post-hoc analysis did not show any effect of the H2 agonist dimaprit (Bonferroni: p>0.999). Instead, infusion of the H3 agonist RAMH significantly increased DA levels, compared to saline (post-hoc Bonferroni, p=0.05). As after HA infusion (Figure 2E-F), there were a small number of animals with very high DA levels following RAMH infusion. This variability in females suggests the possibility of an effect that varies with the estrous cycle.

**Figure 5.**
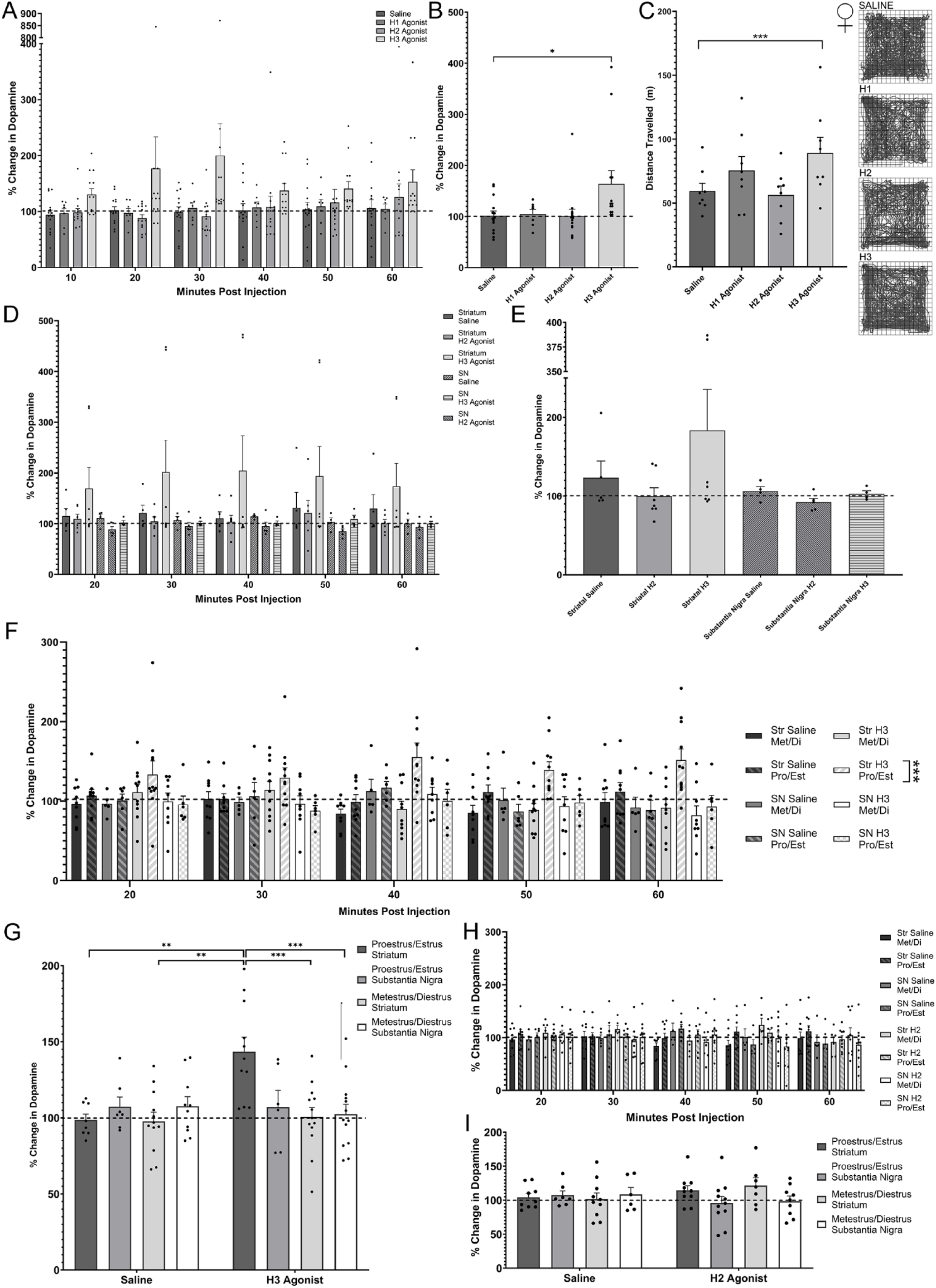
H3R activation in the striatum increases striatal DA in females, but only during the estrus phase of the estrous cycle. **A.** ICV infusion of the H3R agonist RAMH trended towards increased striatal DA in female mice, compared to saline controls and all other treatments when analyzed over the 20-60 minute time interval post-infusion (RM-ANOVA: F[3,44]=2.968, p=0.042; Post-hoc Bonferroni: Saline vs H3, p=0.077). In contrast to males, infusion of H2 agonist dimaprit had no effect (Bonferroni, p>0.999). **B.** Collapsing the data for the 20-60 interval post-infusion to account for individual variance over time shows a significant increase in striatal DA with the infusion of RAMH compared to all other treatments (LMEM: Effect of Treatment, F[3, 44] = 3.417, p=0.025; Post-Hoc Bonferroni (sal vs RAMH), p=0.05). In contrast to what we see in males, there is no effect of dimaprit infusion in females (Bonferroni: p>0.999). **C.** In females, there was a significant increase in locomotion following RAMH infusion compared to saline control (1-way ANOVA: F (1.903, 13.32) = 6.896, p=0.009; Tukey’s (sal vs RAMH): p=0.042, (RAMH vs dimaprit): p=0.027). In contrast to what we see in males, there is no effect of ICV dimaprit infusion [Tukey’s (sal vs H2): p=0.921]. Representative path tracings are shown on the right. **D.** Infusion of RAMH into the striatum appears to increase levels of striatal dopamine over the 20-60 minute interval following infusion, although this did not reach significance (2-Way RM-ANOVA: Treatment x Brain Region F[4,25]=0.740, p=0.487). It appears that this effect is driven by outlier points and causes a bimodal distribution. **E.** When collapsed over the 20-60 minute interval post-infusion, RAMH in the striatum appears to increase levels of striatal dopamine in females, with outlier points still apparent, but fails to reach statistical significance (LMEM: F[6, 25]= 0.792, p=0.464). **F.** When mice are separated into the proestrus/estrus or metestrus/diestrus phases of the estrous cycle, infusion of RAMH into the striatum increases striatal DA only during the proestrus/estrus when measured over the 20-60 minute post infusion interval (3-Way RM-ANOVA: Treatment x Brain Region x Estrus, F[2,87]=2.236, p=0.113, Treatment x Estrus, F[2,87]=2.823, p=0.024, Treatment x Brain Region, F[2,87]=3.436, p=0.037, Brain Region x Estrus, F[1,87]=5.045, p=0.027, Brain Region, F[1,87]=6.507, p=0.012; Fisher’s LSD: P/E Str H3 vs P/E Str Sal, p<0.001, P/E SN H3 vs P/E SN Sal, p=0.698, M/D Str H3 vs M/D Str Sal, p=0.552, M/D SN H3 vs M/D SN Sal, p=0.552). **G.** When collapsed over the 20-60 minute post infusion interval, RAMH in the striatum during pro/estrus remains the only treatment and timing to significantly increase striatal dopamine (LMEM: 3-way interaction, F[12, 48]=2.516, p=0.086; Treatment x Estrus Phase, F[4,48]=3.918, p=0.023, Brain Region x Estrus Phase, F[4,48]=4.094, p=0.046, Treatment x Brain Region, F[4,48]=0.681, p=0.509; Post-Hoc Fisher’s LSD: P/E Str Sal vs P/E Str H3, p=0.079, P/E Str H3 vs P/E SN H3, p=0.003, P/E Str Sal vs M/D Strl Sal p>0.999, P/E SN Sal vs P/E SN H3, p>0.999). **H.** Infusion of dimaprit into the striatum during different estrous phases did not alter striatal dopamine over time (3-Way RM-ANOVA: Fisher’s LSD: P/E Str H2 vs P/E Str Sal, p=0.822, M/D Str H2 vs M/D Str Sal, p=0.240, P/E SN H2 vs P/E SN Sal, p=0.315, M/D SN H2 vs M/D SN Sal, p=0.914). **I.** Collapsed data for infusion of dimaprit into the striatum during different phases of the estrous cycle did not significantly alter levels of striatal DA (LMEM: 3-way interaction N.S.; treatment x brain region: F (1, 61) = 4.377, p=0.041).

We performed a parallel analysis of the effects of histamine agonist infusion on exploratory behavior in the open field. While dimaprit did not cause any behavioral effects compared to saline, RAMH significantly elevated locomotor activity in females, paralleling its effects on DA (Figure 5C; ANOVA: F[1.9,13.3]=6.89, p=0.009, post-hoc Tukey’s H2 p=0.997, H3 p=0.023). When directly comparing behavioral effects across sexes (see Figure 3C; these experiments were run in parallel), sex significantly impacted locomotor behavior (Two-Way ANOVA: no interaction F[3,52]=1.63, p=0.195, effect of sex F[1,52]=56.57, p<0.0001, effect of treatment F[3,52]=3.267, p=0.0285); post hoc analysis (Tukey’s) indicates that females spend more time locomoting compared to males regardless of treatment (p<0.05).

### Experiment Five: Targeted Histamine Agonist Infusion with Estrous Cycle Monitoring in Females. Histamine H3R modulation of DA is regulated by estrous cycle

Targeted infusion of dimaprit and RAMH into the striatum and SNc of females (Figure 1D) did not show a statistically significant effect on dopamine levels over time (Figure 5D, RM-ANOVA: drug x brain region, F[2,25]=0.740, p=0.487; drug F[2,25]=1.018, p=0.376, brain region F[1,25]=1.994, p=0.170) or when collapsed over the 20-60 minutes post infusion (Figure 5E; LMEM: treatment x brain region F[2,25]=0.792, p=0.464, treatment, F[2,25]=1.071 p=0.358, brain region, F[1,25]=1.832 p=0.188). As before (compare Figure 2E,F; Figure 5A-B), several outlier female mice showed very high levels of DA, this time after RAMH infusion into the striatum.

We therefore repeated the experiment with careful monitoring of estrous phase (Figure 1F). This revealed clearer effects (Figure 5F-H). RAMH infusion produced a significant increase in striatal DA, but only in estrus/proestrus, and only when the infusion was into the striatum; there was no effect in metestrus/diestrus, and no effect of RAMH infusion into the SNc. Measurement of DA over time (Figure 5F) showed significant effects of brain region (3-Way RM-ANOVA: brain region F[1,87]=6.507, p=0.012) and interactions between treatment and region (F[2,87]=3.436, p=0.037), treatment and estrous phase (F[2,87]=2.823, p=0.024), and estrous phase and region (brain region x estrous, F[1,87]=5.045, p=0.027; effect of treatment, F[2,87]=2.183, p=0.119, effect of estrous, F[1,87]=1.929, p=0.168, treatment x brain region x estrous, F[2,87]=2.236, p=0.113; Fisher’s LSD: P/E Str H3 vs P/E Str Sal, p<0.001, all other comparisons N.S.). In contrast, infusion of dimaprit did not impact DA levels (Figure 5H, 3-Way Repeated Measure ANOVA, main effects and interactions NS; Fisher’s LSD: P/E Str H2 vs P/E Str Sal, p=0.822, M/D Str H2 vs M/D Str Sal, p=0.240, P/E SN H2 vs P/E SN Sal, p=0.314, M/D SN H2 vs M/D Sal, p=0.914).

To simplify analysis and visualization, we again collapsed the data across 20-60 minutes post-infusion, and confirmed these findings. LMEM analysis shows significant interactions of both treatment and estrous phase (Figure 5G, F[2,97]=3.918, p=0.023) and of brain region and estrous phase (F[1,97]=4.094, p=0.046) with a trend towards a three way interaction (F[2,97]=2.516, p=0.086; treatment x brain region, F[2,97]=0.681, p=0.509, treatment, F[2,97]=0.050, p=0.952, estrous phase, F[1,97]=0.619, p=0.433, brain region, F[1,97]=3.543, p=0.063). Post-hoc analysis showed that infusion of RAMH into the striatum during proestrus/estrus was the only treatment to significantly elevate DA compared to all other conditions (Fisher’s LSD: P/E Str H3 vs P/E Str Sal, p=0.026). Saline or H2 treatments did not differ by region or estrous phase (Figure 5I, Table 2). Thus, activation of H3 receptors in the striatum of females results in significant increases in striatal dopamine only when they are in the proestrus/estrus phase of the estrous cycle. Furthermore, females lacked the H2 response seen in males, regardless of estrous phase.

## Discussion

Here we replicate our previous findings that HA decreases DA levels in the striatum in male mice (Castellan Baldan et al., 2014; Rapanelli et al., 2014) but observe a striking difference in females, in which HA increases striatal DA in some mice. The opposite effects of HA on striatal DA seen in males and females may contribute to some of the sex differences seen in striatal DA dynamics (Zachry et al., 2021), and perhaps to the sex differences seen in the prevalence and presentation of various neuropsychiatric conditions.

Both the striatum and the SN express H1R, H2R, and H3R histamine receptors (Hill and Young, 1980; Martinez-Mir et al., 1990; Vizuete et al., 1997; Pillot et al., 2002; Haas and Panula, 2003; Aceto et al., 2022). In males, we show that activation of H2Rs on GABAergic neurons in the SN mediates HA’s effects. HA has previously been shown to activate these cells *ex vivo* (Korotkova et al., 2002; Zhou et al., 2006). H2R is a G_α_s-coupled receptor that typically excites neurons in which it is expressed (McCormick and Williamson, 1991; Munakata and Akaike, 1994; Haas and Panula, 2003; Haas et al., 2008; Panula and Nuutinen, 2013; Zhuang et al., 2018; Aceto et al., 2022). Our findings *in vivo* suggest that HA activation of H2R on GABAergic interneurons in the SN activates them and thereby inhibits neighboring dopaminergic neurons, inhibiting their firing and thus their release of DA in the striatum – in males. H3R agonists had no significant effect on striatal DA in males.

In females the effect is quite different. Infusion of dimaprit, either ICV or into the SN, does not alter striatal DA. Instead, the H3R agonist RAMH increases striatal DA, and this is mediated in the striatum, not the SN (Figure 5). H3R has a complex role in the striatum, where it functions as an autoreceptor, a presynaptic heteroreceptor, and a postsynaptic receptor (Poli et al., 1991; Schlicker et al., 1994; Pillot et al., 2002; Ellender et al., 2011; González-Sepúlveda et al., 2013; Parks et al., 2014; Bolam and Ellender, 2016; Nieto-Alamilla et al., 2016). Presynaptic H3Rs on DA terminals are unlikely to be responsible for the increase in striatal DA levels, as H3R activity on these terminals would be predicted to decrease DA release (Schlicker et al., 1994). H3 heteroreceptors on GABAergic terminals can suppress GABA release (Schlicker et al., 1994; Li et al., 2014; Takei et al., 2017); RAMH may inhibit GABAergic synapses onto dopaminergic terminals, disinhibiting DA release (Kramer et al., 2020). However, H3R is also expressed postsynaptically on spiny projection neurons (SPNs), the GABAergic principal cells of the striatum, where it interacts with DA receptors and modulates downstream signaling in complex ways (Bolam and Ellender, 2016; Rapanelli et al., 2016; Xu and Pittenger, 2023). H3R on SPNs may modulate their activity and thereby regulate DA release, through either local mechanisms or indirectly, via projections to the SN. Recent work has shown that a subset of D1-expressing SPNs localized to the striosomes directly inhibits SN dopamine neurons (Lazaridis et al., 2024b; Lazaridis et al., 2024a; Okunomiya et al., 2025); inhibition of these neurons could explain the dopamine increase seen after HA infusion in females. These mechanisms remain to be elucidated.

Strikingly, the effects of HA and RAMH in females are modulated by the estrous cycle, with the effects of H3R activation on striatal DA emerging only during proestrus/estrus. This suggests a role for estrogen and/or progesterone, which fluctuate with the estrous cycle, in the modulation of histamine’s effects on dopamine. Estrogen receptors colocalize with HA receptors (Zhou et al., 2007; Kow et al., 2016), and HA-estrogen interactions have been shown to be important in the regulation of hypothalamic neurons (Dupré et al., 2010; Mori et al., 2014). Whether similar interactions occur in the striatum is unclear. If H3-induced inhibition of GABA release, or modulation of SPN activity, is potentiated by sex hormones, then more pronounced functional effects of the H3R agonist RAMH would be expected during phases of the estrous cycle when estrogen and/or progesterone is highest, such as the transition of proestrus into estrus – exactly what we see here. This could also explain the lack of an H3 effect in males. DA levels are known to fluctuate during the estrous cycle, with enhanced DA release during proestrus/estrus (Castner et al., 1993; Xiao and Becker, 1994; Calipari et al., 2017; Yoest et al., 2019; Zachry et al., 2021); these HA-mediated effects may contribute to such fluctuations. While less well characterized than estrogen, progesterone has also been shown to enhance striatal dopamine signaling (Morissette et al., 1990; Yoest et al., 2018). Interestingly, in castrated males treated with estrogen, infusion of progesterone also elevated amphetamine-induced dopamine release (Dluzen and Ramirez, 1990), suggesting that female-specific sex hormones might be working in tandem to impact dopamine release. It will be intriguing to explore whether HA modulation of DA contributes to cyclic changes in neuropsychiatric symptomatology.

Our investigations are motivated by mounting evidence that HA dysregulation contributes to the pathophysiology of TS (Castellan Baldan et al., 2014; Pittenger, 2017). We speculate that the differential effects of HA on striatal DA in males and females may help explain the fact that TS and other tic disorders occur 3-5 more often in males (Comings and Comings, 1985; Freeman et al., 2000; Robertson, 2003; Eapen et al., 2004; Urrutia and Hernández Ch, 2021; Baizabal-Carvallo and Jankovic, 2022; Chou et al., 2022). In males, HA acting on H2R on GABAergic neurons in the SN reduces striatal dopaminergic tone (Figure 3, 4). If this mechanism importantly constrains DA levels at baseline, then reduced HA, such as is in patients or in mice with a TS-associated mutation in the *histidine decarboxylase* gene (Ercan-Sencicek et al., 2010; Castellan Baldan et al., 2014), may disinhibit DA release, leading to the elevation in striatal DA tone that has been inferred in TS patients (Malison et al., 1995; Müller-Vahl et al., 2000; Singer et al., 2002; Serra-Mestres et al., 2004; Wong et al., 2008; Liu et al., 2010; Denys et al., 2013; Hienert et al., 2018) and observed in *Hdc* knockout mice (Castellan Baldan et al., 2014; Rapanelli et al., 2014) If, as our data indicate, this mechanism is absent or attenuated in females, a similar loss of HA may lead to less dysregulation of striatal DA and thus contribute less dramatically to the development of tics. That said, we note that several female members of the originally described family carrying the *HDC* mutation do have TS (Ercan-Sencicek et al., 2010; Castellan Baldan et al., 2014), suggesting that a sufficient reduction in HA is sufficient to cause tics in females.

There is an interesting difference in dopamine dynamics in males and females after saline infusion – dopamine levels rise in males but not in females. These may suggest differences in dopamine response to handling-induced stress. This is consistent with the observation that dopamine rises more in males than in females in response to stress (Dalla et al., 2008).

This study has limitations, of course, and there are areas where further work is needed. The importance of these specific mechanisms in TS and other forms of neuropsychiatric pathophysiology remains to be resolved. The mechanisms whereby estrous phase influences HA’s effects on DA likewise remain to be elucidated, and it is unclear why H2R in the SN does not regulate DA in females. As this is the first time, to our knowledge, that such a marked sexual dimorphism in the effects of HA has been identified, it will be of interest to characterize what other HA-regulated processes also differ in males and females. While we have identified the substantia nigra and striatum as the site of action of histamine in males and females, respectively, other regions may also contribute and will be examined in future experiments. Finally, it will be important to determine whether these effects are recapitulated in humans.

In sum, we have identified unexpected complexity in the regulation of striatal DA by HA, with qualitatively different mechanisms and effects seen in male and female mice. It will be important to further characterize sex differences in the function of the basal ganglia, both in health and in disease states.

## References

Aceto G, Nardella L, Nanni S, Pecci V, Bertozzi A, Colussi C, D’Ascenzo M, Grassi C (2022) Activation of histamine type 2 receptors enhances intrinsic excitability of medium spiny neurons in the nucleus accumbens. J Physiol 600:2225–2243.

Airaksinen MS, Panula P (1988) The histaminergic system in the guinea pig central nervous system: an immunocytochemical mapping study using an antiserum against histamine. J Comp Neurol 273:163–186.

Almey A, Milner TA, Brake WG (2015) Estrogen receptors in the central nervous system and their implication for dopamine-dependent cognition in females. Horm Behav 74:125–138.

Andersen ML, Sawyer EK, Howell LL (2012) Contributions of neuroimaging to understanding sex differences in cocaine abuse. Exp Clin Psychopharmacol 20:2–15.

Anichtchik OV, Rinne JO, Kalimo H, Panula P (2000) An altered histaminergic innervation of the substantia nigra in Parkinson’s disease. Exp Neurol 163:20–30.

Arrang JM, Garbarg M, Lancelot JC, Lecomte JM, Pollard H, Robba M, Schunack W, Schwartz JC (1987) Highly potent and selective ligands for histamine H3-receptors. Nature 327:117–123.

Arvidsson E, Viereckel T, Mikulovic S, Wallen-Mackenzie A (2014) Age- and sex-dependence of dopamine release and capacity for recovery identified in the dorsal striatum of C57/Bl6J mice. PLoS One 9:e99592.

Baizabal-Carvallo JF, Jankovic J (2022) Sex differences in patients with Tourette syndrome. CNS Spectr:1–7.

Bazzett TJ, Becker JB (1994) Sex differences in the rapid and acute effects of estrogen on striatal D2 dopamine receptor binding. Brain Research 637:163–172.

Becker JB (1990) Direct effect of 17β-estradiol on striatum: Sex differences in dopamine release. Synapse 5:157–164.

Becker JB (1999) Gender Differences in Dopaminergic Function in Striatum and Nucleus Accumbens. Pharmacology Biochemistry and Behavior 64:803–812.

Bolam JP, Ellender TJ (2016) Histamine and the striatum. Neuropharmacology 106:74–84.

Byers SL, Wiles MV, Dunn SL, Taft RA (2012) Mouse Estrous Cycle Identification Tool and Images. PLOS ONE 7:e35538.

Calipari ES, Juarez B, Morel C, Walker DM, Cahill ME, Ribeiro E, Roman-Ortiz C, Ramakrishnan C, Deisseroth K, Han MH, Nestler EJ (2017) Dopaminergic dynamics underlying sex-specific cocaine reward. Nat Commun 8:13877.

Castellan Baldan L et al. (2014) Histidine decarboxylase deficiency causes tourette syndrome: parallel findings in humans and mice. Neuron 81:77--90.

Castner SA, Xiao L, Becker JB (1993) Sex differences in striatal dopamine: in vivo microdialysis and behavioral studies. Brain Research 610:127–134.

Chou IJ, Hung PC, Lin JJ, Hsieh MY, Wang YS, Kuo CY, Kuo CF, Lin KL, Wang HS (2022) Incidence and prevalence of Tourette syndrome and chronic tic disorders in Taiwan: a nationwide population-based study. Soc Psychiatry Psychiatr Epidemiol 57:1711–1721.

Comings DE, Comings BG (1985) Tourette syndrome: clinical and psychological aspects of 250 cases. Am J Hum Genet 37:435–450.

Creutz LM, Kritzer MF (2004) Mesostriatal and mesolimbic projections of midbrain neurons immunoreactive for estrogen receptor beta or androgen receptors in rats. J Comp Neurol 476:348–362.

Dalla C, Antoniou K, Kokras N, Drossopoulou G, Papathanasiou G, Bekris S, Daskas S, Papadopoulou-Daifoti Z (2008) Sex differences in the effects of two stress paradigms on dopaminergic neurotransmission. Physiology & Behavior 93:595–605.

Denys D, de Vries F, Cath D, Figee M, Vulink N, Veltman DJ, van der Doef TF, Boellaard R, Westenberg H, van Balkom A, Lammertsma AA, van Berckel BNM (2013) Dopaminergic activity in Tourette syndrome and obsessive-compulsive disorder. European Neuropsychopharmacology 23:1423–1431.

Dluzen DE, Ramirez VD (1990) In vitro progesterone modulates amphetamine-stimulated dopamine release from the corpus striatum of castrated male rats treated with estrogen. Neuroendocrinology 52:517–520.

Dupré C, Lovett-Barron M, Pfaff DW, Kow LM (2010) Histaminergic responses by hypothalamic neurons that regulate lordosis and their modulation by estradiol. Proc Natl Acad Sci U S A 107:12311–12316.

Durant GJ, Ganellin CR, Parsons ME (1977) Dimaprit, [S-[3-(N,N-dimethylamino)propyl]isothiourea]. A highly specific histamine H2-receptor agonist. Part 2. Structure-activity considerations. Agents and Actions 7:39–43.

Eapen V, Fox-Hiley P, Banerjee S, Robertson M (2004) Clinical features and associated psychopathology in a Tourette syndrome cohort. Acta Neurol Scand 109:255–260.

Ellender TJ, Huerta-Ocampo I, Deisseroth K, Capogna M, Bolam JP (2011) Differential Modulation of Excitatory and Inhibitory Striatal Synaptic Transmission by Histamine. The Journal of Neuroscience 31:15340–15351.

Ercan-Sencicek GA et al. (2010) L-Histidine Decarboxylase and Tourette’s Syndrome. The New England Journal of Medicine 362:1901–1908.

Franklin KBJ, Paxinos G (1997) The mouse brain in stereotaxic coordinates. San Diego: Academic Press San Diego.

Freeman RD, Fast DK, Burd L, child… K-Ja (2000) An international perspective on Tourette syndrome: selected findings from 3500 individuals in 22 countries.

Garcia-Delgar B, Servera M, Coffey BJ, Lázaro L, Openneer T, Benaroya-Milshtein N, Steinberg T, Hoekstra PJ, Dietrich A, Morer A (2022) Tic disorders in children and adolescents: does the clinical presentation differ in males and females? A report by the EMTICS group. Eur Child Adolesc Psychiatry 31:1539–1548.

González-Sepúlveda M, Rosell S, Hoffmann HM, Castillo-Ruiz MdM, Mignon V, Moreno-Delgado D, Vignes M, Díaz J, Sabriá J, Ortiz J (2013) Cellular distribution of the histamine H3 receptor in the basal ganglia: Functional modulation of dopamine and glutamate neurotransmission. Basal Ganglia 3:109–121.

Haas H, Panula P (2003) The role of histamine and the tuberomamillary nucleus in the nervous system. Nature Reviews Neuroscience 4:121–130.

Haas HL, Sergeeva OA, Selbach O (2008) Histamine in the nervous system. Physiol Rev 88:1183–1241.

Hahn A, Reed MB, Pichler V, Michenthaler P, Rischka L, Godbersen GM, Wadsak W, Hacker M, Lanzenberger R (2021) Functional dynamics of dopamine synthesis during monetary reward and punishment processing. J Cereb Blood Flow Metab 41:2973–2985.

Hienert M, Gryglewski G, Stamenkovic M, Kasper S, Lanzenberger R (2018) Striatal dopaminergic alterations in Tourette’s syndrome: a meta-analysis based on 16 PET and SPECT neuroimaging studies. Translational Psychiatry 8:143.

Hill SJ, Young JM (1980) Histamine H1-receptors in the brain of the guinea-pig and the rat: differences in ligand binding properties and regional distribution. Br J Pharmacol 68:687–696.

Honrubia MA, Vilaró MT, Palacios JM, Mengod G (2000) Distribution of the histamine H(2) receptor in monkey brain and its mRNA localization in monkey and human brain. Synapse 38:343–354.

Kaasinen V, Någren K, Hietala J, Farde L, Rinne JO (2001) Sex Differences in Extrastriatal Dopamine D2-Like Receptors in the Human Brain. American Journal of Psychiatry 158:308–311.

Korotkova TM, Haas HL, Brown RE (2002) Histamine excites GABAergic cells in the rat substantia nigra and ventral tegmental area in vitro. Neuroscience Letters 320:133–136.

Kow LM, Pataky S, Dupré C, Phan A, Martin-Alguacil N, Pfaff DW (2016) Analyses of rapid estrogen actions on rat ventromedial hypothalamic neurons. Steroids 111:100–112.

Kramer PF, Twedell EL, Shin JH, Zhang R, Khaliq ZM (2020) Axonal mechanisms mediating γ-aminobutyric acid receptor type A (GABA-A) inhibition of striatal dopamine release. eLife 9:e55729, citation = eLife 52020;55729 e55729.

Krause M, Rouleau A, Stark H, Luger P, Lipp R, Garbarg M, Schwart JC, Schunack W (1995) Synthesis, X-ray crystallography, and pharmacokinetics of novel azomethine prodrugs of (R)-alpha-methylhistamine: highly potent and selective histamine H3 receptor agonists. J Med Chem 38:4070–4079.

Laakso A, Vilkman H, Bergman Jö, Haaparanta M, Solin O, Syvälahti E, Salokangas RKR, Hietala J (2002) Sex differences in striatal presynaptic dopamine synthesis capacity in healthy subjects. Biological Psychiatry 52:759–763.

Larsh TR, Wu SW, Huddleston DA, White S, Lipps TD, Gilbert DL (2023) Adolescent Gender Differences in Tic- and Non-Tic-Related Impairments in Tourette Syndrome. J Child Neurol 38:283–289.

Lazaridis I, Crittenden JR, Ahn G, Hirokane K, Yoshida T, Mahar A, Skara V, Meletis K, Parvataneni K, Ting JT, Hueske E, Matsushima A, Graybiel AM (2024a) Striosomes Target Nigral Dopamine-Containing Neurons via Direct-D1 and Indirect-D2 Pathways Paralleling Classic Direct-Indirect Basal Ganglia Systems. bioRxiv:2024.2006.2001.596922.

Lazaridis I, Crittenden JR, Ahn G, Hirokane K, Wickersham IR, Yoshida T, Mahar A, Skara V, Loftus JH, Parvataneni K, Meletis K, Ting JT, Hueske E, Matsushima A, Graybiel AM (2024b) Striosomes control dopamine via dual pathways paralleling canonical basal ganglia circuits. Current Biology 34:5263–5283.e5268.

Li WW, Cheng LZ, Zou Z, Tian ML, Zhang H, Raya AD, Wang Y, Shi XY (2014) (R)-alpha-methylhistamine suppresses inhibitory neurotransmission in hippocampal CA1 pyramidal neurons counteracting propofol-induced amnesia in rats. CNS Neurosci Ther 20:851–859.

Liu H, Dong F, Meng Z, Zhang B, Tan J, Wang Y (2010) Evaluation of Tourette’s syndrome by 99mTc-TRODAT-1 SPECT/CT imaging. Annals of Nuclear Medicine 24:515–521.

Malén T, Karjalainen T, Isojärvi J, Vehtari A, Bürkner PC, Putkinen V, Kaasinen V, Hietala J, Nuutila P, Rinne J, Nummenmaa L (2022) Atlas of type 2 dopamine receptors in the human brain: Age and sex dependent variability in a large PET cohort. Neuroimage 255:119149.

Malison RT, McDougle CJ, Dyck CHv, Scahill L, Baldwin RM, Seibyl JP, Price LH, Leckman JF, Innis RB (1995) [123I]beta-CIT SPECT imaging of striatal dopamine transporter binding in Tourette’s disorder. The American journal of psychiatry 152:1359–1361.

Malmberg-Aiello P, Lamberti C, Ipponi A, Bartolini A, Schunack W (1998) Evidence for hypernociception induction following histamine H1 receptor activation in rodents. Life Sci 63:463–476.

Manza P, Shokri-Kojori E, Wiers CE, Kroll D, Feldman D, McPherson K, Biesecker E, Dennis E, Johnson A, Kelleher A, Qu S, Tomasi D, Wang G-J, Volkow ND (2022) Sex differences in methylphenidate-induced dopamine increases in ventral striatum. Molecular Psychiatry 27:939–946.

Martinez-Mir MI, Pollard H, Moreau J, Arrang JM, Ruat M, Traiffort E, Schwartz JC, Palacios JM (1990) Three histamine receptors (H1, H2 and H3) visualized in the brain of human and non-human primates. Brain Research 526:322–327.

McCormick DA, Williamson A (1991) Modulation of neuronal firing mode in cat and guinea pig LGNd by histamine: possible cellular mechanisms of histaminergic control of arousal. J Neurosci 11:3188–3199.

McDermott JL, Dluzen DE (2007) Aging and sex differences in striatal dopaminergic function. Neuroscience 149:401–408.

McDermott JL, Liu B, Dluzent DE (1994) Sex Differences and Effects of Estrogen on Dopamine and DOPAC Release from the Striatum of Male and Female CD-1 Mice. Experimental Neurology 125:306–311.

Michinaga S, Sonoda K, Inazuki N, Ezaki M, Awane H, Shimizu K, Hishinuma S, Mizuguchi H (2022) Selective histamine H2 receptor agonists alleviate blood-brain barrier disruption by promoting the expression of vascular protective factors following traumatic brain injury in mice. Journal of Pharmacological Sciences 150:135–145.

Mori H, Matsuda K, Yamawaki M, Kawata M (2014) Estrogenic regulation of histamine receptor subtype H1 expression in the ventromedial nucleus of the hypothalamus in female rats. PLoS One 9:e96232.

Morissette M, Lévesque D, Bélanger A, Paolo TD (1990) A physiological dose of estradiol with progesterone affects striatum biogenic amines. Canadian Journal of Physiology and Pharmacology 68:1520–1526.

Müller-Vahl KR, Berding G, Brücke T, Kolbe H, Meyer GJ, Hundeshagen H, Dengler R, Knapp WH, Emrich HM (2000) Dopamine transporter binding in Gilles de la Tourette syndrome. Journal of Neurology 247:514–520.

Munakata M, Akaike N (1994) Regulation of K+ conductance by histamine H1 and H2 receptors in neurones dissociated from rat neostriatum. J Physiol 480 (Pt 2):233–245.

Munro CA, McCaul ME, Wong DF, Oswald LM, Zhou Y, Brasic J, Kuwabara H, Kumar A, Alexander M, Ye W, Wand GS (2006) Sex Differences in Striatal Dopamine Release in Healthy Adults. Biological Psychiatry 59:966–974.

Nieto-Alamilla G, Márquez-Gómez R, García-Gálvez AM, Morales-Figueroa GE, Arias-Montaño JA (2016) The Histamine H3 Receptor: Structure, Pharmacology, and Function. Mol Pharmacol 90:649–673.

Oishi R, Itoh Y, Nishibori M, Saeki K (1989) Effects of the histamine H3-agonist (R)-alpha-methylhistamine and the antagonist thioperamide on histamine metabolism in the mouse and rat brain. J Neurochem 52:1388–1392.

Okunomiya T, Watanabe D, Banno H, Kondo T, Imamura K, Takahashi R, Inoue H (2025) Striosome Circuitry Stimulation Inhibits Striatal Dopamine Release and Locomotion. J Neurosci 45.

Ovtscharoff W, Eusterschulte B, Zienecker R, Reisert I, Pilgrim C (1992) Sex differences in densities of dopaminergic fibers and GABAergic neurons in the prenatal rat striatum. Journal of Comparative Neurology 323:299–304.

Panula P, Nuutinen S (2013) The histaminergic network in the brain: basic organization and role in disease. Nat Rev Neurosci 14:472–487.

Parks GS, Olivas ND, Ikrar T, Sanathara NM, Wang L, Wang Z, Civelli O, Xu X (2014) Histamine inhibits the melanin-concentrating hormone system: implications for sleep and arousal. The Journal of Physiology 592:2183–2196.

Patel D, Vishwakarma PK, Patel R, Jain NS (2021) Central histaminergic transmission modulates the expression of chronic nicotine withdrawal induced anxiety-like and somatic behavior in mice. Behav Brain Res 399:112997.

Pillot C, Heron A, Cochois V, Tardivel-Lacombe J, Ligneau X, Schwartz JC, Arrang JM (2002) A detailed mapping of the histamine H(3) receptor and its gene transcripts in rat brain. Neuroscience 114:173–193.

Pittenger C (2017) Histidine Decarboxylase Knockout Mice as a Model of the Pathophysiology of Tourette Syndrome and Related Conditions. Handb Exp Pharmacol 241:189–215.

Pittenger C (2020) The histidine decarboxylase model of tic pathophysiology: a new focus on the histamine H(3) receptor. Br J Pharmacol 177:570–579.

Pittenger C (2024) The Neurobiology of Tic Disorders and Obsessive-Compulsive Disorder: Animal and Human Studies. In: Charney & Nestler’s Neurobiology of Mental Illness (Charney D, Nestler, E., Buxbaum, J.D., Binder, E.B., Gordon, J.A., and Picciotto, M.R., ed). New York: Oxford University Press.

Poli E, Coruzzi G, Bertaccini G (1991) Histamine H3 receptors regulate acetylcholine release from the guinea pig ileum myenteric plexus. Life Sci 48:Pl63–68.

Purón-Sierra L, Miranda MI (2014) Histaminergic modulation of cholinergic release from the nucleus basalis magnocellularis into insular cortex during taste aversive memory formation. PLoS One 9:e91120.

Rapanelli M, Frick L, Bito H, Pittenger C (2017a) Histamine modulation of the basal ganglia circuitry in the development of pathological grooming. Proc Natl Acad Sci U S A 114:6599–6604.

Rapanelli M, Frick L, Pogorelov V, Ohtsu H, Bito H, Pittenger C (2017b) Histamine H3R receptor activation in the dorsal striatum triggers stereotypies in a mouse model of tic disorders. Transl Psychiatry 7:e1013.

Rapanelli M, Frick LR, Pogorelov V, Ota KT, Abbasi E, Ohtsu H, Pittenger C (2014) Dysregulated intracellular signaling in the striatum in a pathophysiologically grounded model of Tourette syndrome. European Neuropsychopharmacology 24:1896--1906.

Rapanelli M, Frick LR, Horn KD, Schwarcz RC, Pogorelov V, Nairn AC, Pittenger C (2016) The Histamine H3 Receptor Differentially Modulates Mitogen-activated Protein Kinase (MAPK) and Akt Signaling in Striatonigral and Striatopallidal Neurons. J Biol Chem 291:21042–21052.

Robertson MM (2003) Diagnosing Tourette syndrome: is it a common disorder? J Psychosom Res 55:3–6.

Savage JT, Ramirez JJ, Risher WC, Wang Y, Irala D, Eroglu C (2024) SynBot is an open-source image analysis software for automated quantification of synapses. Cell Reports Methods 4:100861.

Schindelin J, Arganda-Carreras I, Frise E, Kaynig V, Longair M, Pietzsch T, Preibisch S, Rueden C, Saalfeld S, Schmid B, Tinevez J-Y, White DJ, Hartenstein V, Eliceiri K, Tomancak P, Cardona A (2012) Fiji: an open-source platform for biological-image analysis. Nature Methods 9:676–682.

Schlicker E, Malinowska B, Kathmann M, Göthert M (1994) Modulation of neurotransmitter release via histamine H3 heteroreceptors. Fundam Clin Pharmacol 8:128–137.

Schwabe MJ, Konkol RJ (1992) Menstrual cycle-related fluctuations of tics in Tourette syndrome. Pediatr Neurol 8:43–46.

Serra-Mestres J, Ring HA, Costa DC, Gacinovic S, Walker Z, Lees AJ, Robertson MM, Trimble MR (2004) Dopamine transporter binding in Gilles de la Tourette syndrome: a [123I]FP-CIT/SPECT study. Acta Psychiatrica Scandinavica 109:140–146.

Singer HS, Szymanski S, Giuliano J, Yokoi F, Dogan AS, Brasic JR, Zhou Y, Grace AA, Wong DF (2002) Elevated Intrasynaptic Dopamine Release in Tourette’s Syndrome Measured by PET. American Journal of Psychiatry 159:1329–1336.

Takagi H, Morishima Y, Matsuyama T, Hayashi H, Watanabe T, Wada H (1986) Histaminergic axons in the neostriatum and cerebral cortex of the rat: a correlated light and electron microscopic immunocytochemical study using histidine decarboxylase as a marker. Brain Res 364:114–123.

Takei H, Yamamoto K, Bae YC, Shirakawa T, Kobayashi M (2017) Histamine H(3) Heteroreceptors Suppress Glutamatergic and GABAergic Synaptic Transmission in the Rat Insular Cortex. Front Neural Circuits 11:85.

Urrutia SV, Hernández Ch M (2021) [A pediatric cohort with Gilles de la Tourette syndrome]. Andes Pediatr 92:838–846.

Vizuete ML, Traiffort E, Bouthenet ML, Ruat M, Souil E, Tardivel-Lacombe J, Schwartz JC (1997) Detailed mapping of the histamine H2 receptor and its gene transcripts in guinea-pig brain. Neuroscience 80:321–343.

Vohora D, Pal SN, Pillai KK (2001) Histamine and selective H3-receptor ligands: a possible role in the mechanism and management of epilepsy. Pharmacology Biochemistry and Behavior 68:735–741.

Walker QD, Ray R, Kuhn CM (2006) Sex Differences in Neurochemical Effects of Dopaminergic Drugs in Rat Striatum. Neuropsychopharmacology 31:1193–1202.

Walker QD, Rooney MB, Wightman RM, Kuhn CM (2000) Dopamine release and uptake are greater in female than male rat striatum as measured by fast cyclic voltammetry. Neuroscience 95:1061–1070.

Wong DF, Brašić JR, Singer HS, Schretlen DJ, Kuwabara H, Zhou Y, Nandi A, Maris MA, Alexander M, Ye W, Rousset O, Kumar A, Szabo Z, Gjedde A, Grace AA (2008) Mechanisms of Dopaminergic and Serotonergic Neurotransmission in Tourette Syndrome: Clues from an In Vivo Neurochemistry Study with PET. Neuropsychopharmacology 33:1239–1251.

Xiao L, Becker JB (1994) Quantitative microdialysis determination of extracellular striatal dopamine concentration in male and female rats: effects of estrous cycle and gonadectomy. Neurosci Lett 180:155–158.

Xu J, Pittenger C (2023) The histamine H3 receptor modulates dopamine D2 receptor-dependent signaling pathways and mouse behaviors. J Biol Chem 299:104583.

Yoest KE, Quigley JA, Becker JB (2018) Rapid effects of ovarian hormones in dorsal striatum and nucleus accumbens. Horm Behav 104:119–129.

Yoest KE, Cummings JA, Becker JB (2019) Oestradiol influences on dopamine release from the nucleus accumbens shell: sex differences and the role of selective oestradiol receptor subtypes. Br J Pharmacol 176:4136–4148.

Yu L, Liao PC (2000) Sexual differences and estrous cycle in methamphetamine-induced dopamine and serotonin depletions in the striatum of mice. Journal of Neural Transmission 107:419–427.

Zachry JE, Nolan SO, Brady LJ, Kelly SJ, Siciliano CA, Calipari ES (2021) Sex differences in dopamine release regulation in the striatum. Neuropsychopharmacology 46:491–499.

Zhou FW, Xu JJ, Zhao Y, LeDoux MS, Zhou FM (2006) Opposite functions of histamine H1 and H2 receptors and H3 receptor in substantia nigra pars reticulata. J Neurophysiol 96:1581–1591.

Zhou J, Lee AW, Devidze N, Zhang Q, Kow LM, Pfaff DW (2007) Histamine-induced excitatory responses in mouse ventromedial hypothalamic neurons: ionic mechanisms and estrogenic regulation. J Neurophysiol 98:3143–3152.

Zhuang QX, Xu HT, Lu XJ, Li B, Yung WH, Wang JJ, Zhu JN (2018) Histamine Excites Striatal Dopamine D1 and D2 Receptor-Expressing Neurons via Postsynaptic H1 and H2 Receptors. Mol Neurobiol 55:8059–8070.

